# Single-cell RNA sequencing reveals novel cell differentiation dynamics during human airway epithelium regeneration

**DOI:** 10.1101/451807

**Authors:** Sandra Ruiz Garcia, Marie Deprez, Kevin Lebrigand, Agnès Paquet, Amélie Cavard, Marie-Jeanne Arguel, Virginie Magnone, Ignacio Caballero, Sylvie Leroy, Charles-Hugo Marquette, Brice Marcet, Pascal Barbry, Laure-Emmanuelle Zaragosi

## Abstract

**Background:** It is usually considered that the upper airway epithelium is composed of multiciliated, goblet, secretory and basal cells, which collectively constitute an efficient first line of defense against inhalation of noxious substances. Upon injury, regeneration of this epithelium through proliferation and differentiation can restore a proper mucociliary function. However, in chronic airway diseases, the injured epithelium frequently displays defective repair leading to tissue remodeling, characterized by a loss of multiciliated cells and mucus hyper-secretion. Delineating drivers of differentiation dynamics and cell fate in the human airway epithelium is important to preserve homeostasis.

**Results:** We have used single cell transcriptomics to characterize the sequence of cellular and molecular processes taking place during human airway epithelium regeneration. We have characterized airway subpopulations with high resolution and lineage inference algorithms have unraveled cell trajectories from basal to luminal cells, providing markers for specific cell populations, such as deuterosomal cells, i.e. precursors of multiciliated cells. We report that goblet cells, like secretory cells, can act as precursors of multiciliated cells. Our study provides a repertoire of molecules involved in key steps of the regeneration process, either keratins or components of the Notch, Wnt or BMP/TGFβ signaling pathways. Our findings were confirmed in independent experiments performed on fresh human and pig airway samples, and on mouse tracheal epithelial cells.

**Conclusions:** Our single-cell RNA-seq study provides novel insights about airway epithelium differentiation dynamics, clarifies cell trajectories between secretory, goblet and multiciliated cells, identifies novel cell subpopulations, and maps the activation and repression of key signaling pathways.

## Background

In mammalian airways, a pseudo-stratified mucociliary epithelium constitutes an efficient first line of defense of the respiratory tract against a large panel of inhaled substances. This epithelium forms a complex ecosystem, mainly composed by: multiciliated cells (MCCs), projecting hundreds of motile cilia at their apical surface, goblet cells (GCs), secreting protective mucins on the luminal surface, secretory cells (SCs), producing anti-microbial and anti-inflammatory peptides, and basal cells, playing a role in adhesion and stability of the epithelium [1, 2]. Altered balance between multiciliated and goblet lineages (i.e. decreased number of MCCs with increased number of GCs) is a hallmark of chronic respiratory diseases, such as chronic obstructive pulmonary disease, primary ciliary dyskinesia, asthma or cystic fibrosis. These diseases, which collectively affect hundreds of millions of people, are characterized by frequent injuries, repair defects, tissue remodeling and altered mucociliary clearance [3-5]. In order to accurately characterize the cellular and molecular identity of progenitor cells of the airway epithelium that contribute to efficient tissue regeneration and of the mechanisms that control the MCC/GC balance, key facts about lung epithelial cell fates have been investigated in mouse, mainly injuring the epithelium with noxious agents. Lineage tracing studies have identified basal cells as the main airway progenitor cells, as they display self-renewal capacities and ability to differentiate into MCCs and SCs [2, 6, 7]. Basal cells in mouse are abundant in the trachea and in the main bronchi, but they are absent in smaller airways [8]. In human, they populate the whole airways, though showing a 2-fold decrease in numbers in smaller airways, i.e. with diameters below 0.5mm [9]. The TP63 transcription factor, keratins 5 and 14 (KRT5 and KRT14), podoplanin (PDPN), nerve growth factor receptor (NGFR), galectin 1 (LGALS1), integrins α6 and β4 (ITGA6 and ITGB4), laminins α3 and β3 (LAMA3 and LAMB3) have been frequently used as selective markers of BCs [2, 6]. While a direct differentiation of a subpopulation of BCs into MCCs has been reported upon injury [10], the consensus emerging from lineage tracing studies suggests that BCs usually differentiate first into SCs [11]. SCs, known as club (or Clara) cells, are widespread from mouse trachea to bronchioles, and are also found in the human airways, though at a much less abundance, being absent from the upper airways and enriched in the terminal and respiratory bronchioles [12]. SCs are detected at a luminal position of the epithelium, showing a characteristic columnar shape, being filled with secretory granules containing anti-microbial, anti-inflammatory peptides [13] and contributing to xenobiotic metabolism [14]. The most extensively studied marker of SCs is the anti-inflammatory secretoglobin SCGB1A1. Lineage tracing studies have used SCGB1A1 to determine the fate of SCs and have shown that they give rise to MCCs, identified by the specific expression of the transcription factor FOXJ1 [11, 15] and to GCs, identified by the expression of specific mucin MUC5AC [2, 16]. Molecular mechanisms regulating cell fate decisions in the airway epithelium lineages have been investigated over the past few years and they have established the role of Notch signaling during commitment of BCs. It is now clear that Notch activation leads to the SC/GC lineages as Notch inhibition leads to commitment towards the MCC fate [17-20]. In agreement with these findings, our group has shown that Notch pathway inhibition by the miR-34/449 families of microRNAs is required for MCC differentiation [21, 22]. Although *in vivo* lineage tracing studies have largely unveiled cell lineage hierarchies in the airway epithelium, they have some inherent limitations. They can only be performed in mouse, after specific, drastic, non-physiological forms of injuries which may not completely reveal physiological tissue turnover. Moreover, they rely on genetic cell labelling, usually *Krt5* for BCs and *Scgb1a1* for SCs, thus, they are not necessarily comprehensive and cannot provide a full picture of cell hierarchies in the airway epithelium. In human, there is no possibility to perform such studies, and cell lineage hierarchies must be inferred indirectly by *in vitro* approaches. Single-cell RNA-sequencing, coupled with computational methods measuring transition between cell states, has emerged as a complementary and powerful approach to reveal lineage hierarchies in many tissues [23-25] and even in a whole organism, by capturing undifferentiated, intermediate and terminally differentiated states [26]. In the lung, one pioneer single-cell study on 198 cells has delineated lineage hierarchies of alveolar cells in the mouse distal lung [27]. Recent studies have established first atlases of the airways in mouse [28] and human [29, 30]. However, extensive characterization of cell population diversity and cell lineages in the human airways remains to be achieved. Here, we have performed and collected single-cell RNA-seq data from fresh human airway epithelial tissues and all along an experiment of 3D *in vitro* regeneration of human airway epithelium. Our work has resulted in a comprehensive cell trajectory roadmap of human airways, which identifies novel cell populations and offer new insights into molecular mechanisms taking place during the mucociliary epithelium regeneration.

## Results

### Reconstruction of cell lineage in regenerating airway epithelium by single-cell RNA-seq

To identify trajectories of cells composing the human airway epithelium, we have analyzed single-cell transcriptomes obtained at successive stages during in vitro 3D regeneration of this tissue (Fig. 1a, b). We first validated that this *in vitro* model faithfully recapitulated native airway tissues by comparing cell population compositions of 3D regenerated airway tissues (Human Airway Epithelial Cells, HAECs) with that of fresh human airway tissues. We performed single-cell RNA-seq of epithelial cells dissociated from nasal brushing samples and from nasal turbinates which were resected during surgery, as well as single-cell RNA-seq of HAECs at late time point of *in vitro* air-liquid interface differentiation (3D cells). Our main results were obtained with HAECs that were cultured in a Pneumacult media (StemCell Technologies), which allows the production of multiciliated and goblet cells, but additional experiments were also performed with HAECs cultured in BEGM (Lonza), which rather favors the production of multiciliated cells. Cell identity was inferred from the expression of specific marker genes, such as *KRT5* and *TP63* for basal cells (BCs), *SCGB1A1* for secretory cells (SCs), *MUC5AC* for goblet cells (GCs) and *FOXJ1* for multiciliated cells (MCCs). These cell types were robustly found in all samples at various proportions, validating the use of these *in vitro* models to trace airway cell lineages (Supplemental Figure S1a-c). We also confirmed that cell type proportions inferred from single-cell RNA-seq were correlated with cell type proportions inferred from protein measurements by performing immunostaining of selected population markers (Supplemental Figure S1d, e). In addition, we have evaluated the effect of cell dissociation on gene expression (Supplemental Figure S2a). We found that cell dissociation did not produce a major impact on gene expression with the exception of *FOS* and *FOSB* which were highly upregulated (Supplemental Figure S2b, c). Molecular function enrichment with Ingenuity Pathway Analysis (Qiagen) showed that “cell death and survival” and “cellular growth and proliferation” were the only molecular functions to be regulated with a p-value inferior to 0.001 (Supplemental Figure S2d).

**Figure 1:**
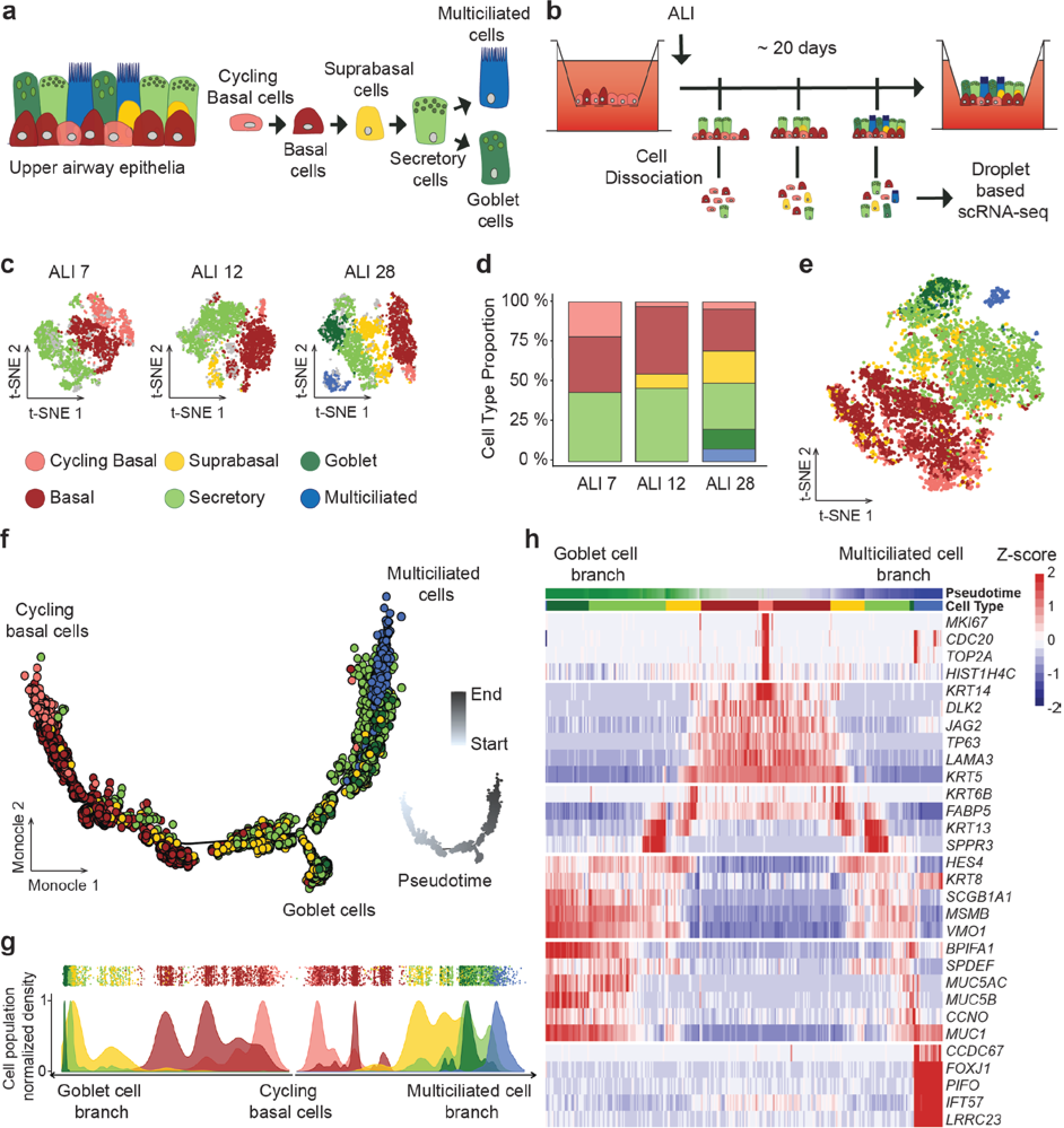
Characterization of MCC and Goblet cell lineages during airway epithelium regeneration using single cell RNA-Seq. **a** Model of upper airway epithelium, based on 6 major types of epithelial cells, with consensus lineage hierarchy. **b** single-cell (sc)RNA-seq experimental design. Regenerating airway epithelia was dissociated at successive days (7, 12, 28) after a transition to an air-liquid interface (ALI). **c** t-SNE plots of the scRNA-seq expression data highlighting the main cell types observed at ALI7 (3426 cells), ALI12 (2785 cells), ALI28 (3615 cells) (grey: unassigned cells). **d** Relative abundance of the 6 main cell types at each time point. **e** Aggregate t-SNE plot of gene expression in 9826 cells. **f** Inference of goblet and MCC cell lineages by Monocle 2, based on an aggregate of the entire experiment. Color code is the same as in c. Inset: pseudotime picturing by a white to grey gradient along the differentiation trajectory. **g** Distribution of the 6 main cell types in the pseudotime along the two branches of the trajectory (down: goblet cell branch; upper right: multiciliated cell branch). **h** Heatmap representing the smoothened temporal expression pattern of a representative list of cell type specific markers, with branch representation as in **g**. Cells were ordered by branch, then cluster appearance, then pseudotime.

We have generated single-cell transcriptomes of HAECs cultivated in Pneumacult medium at 3 time points of *in vitro* airway regeneration (ALI7, ALI12, and ALI28) (Fig. 1b). Time points were chosen as most representative of the known steps of airway regeneration: proliferation, polarization and specification [31]. This experiment was complemented by 6 additional time points of HAECs cultivated in BEGM medium (ALI2, ALI4, ALI7, ALI12, ALI17 and ALI22). In a first approach, each time point was analyzed independently. We did 10 random selections of cells, corresponding to subgroups containing 90% of the initial number of cells. The resulting gene expression submatrices were then iteratively clustered (10 times with varying parameters), and a census was applied to define the most robust cell types. We then studied the variations of these populations during the entire time course. Cells clustered in 6 main populations in the Pneumacult condition: (1) cycling *(MKI67*+) and (2) non-cycling (*MKI67*-) BCs (*KRT5*+/*TP63*+), (3) supraBCs (*KRT5*+/ *TP63*-/*KRT13*+/*KRT4*+), (4) SCs (*SCGB1A1*+), (5) GCs (*MUC5AC*+) and (6) MCCs (*FOXJ1*+) (Figure 1c). Cell population proportions evolved during the time course, with a global reduction of BCs and SCs, a first detection of supraBCs at ALI7, followed by an increase of the proportion of this cell population at ALI28 and a first detection of GCs and MCCs at ALI28 (Fig. 1d). In the BEGM condition, cells clustered in 7 cell populations (Supplemental Figure S3a, b). We did not detect SCs and GCs with this culture condition, but we found instead a cell population that we termed “Secretory-like cells”, given their high gene expression similarity with SCs, except for *SCGB1A1*, which was not detected (Supplemental Figure S4). Additional cell types were found in these samples: *KRT5*-supraBCs (*TP63*-/*KRT13*+/*KRT4*+) and 2 cell populations that we termed as “undefined intermediates 1” and “undefined intermediates 2” because their gene expression profile did not allow unambiguous classification. Differential gene expression showed that these undefined intermediates 1 and 2 expressed specific genes, such as *SPINK1* and *SPINK7*, respectively. A representation of all cells analyzed in each culture was generated using t-SNE graphs on the aggregate of all time points for each medium condition (Fig. 1e for Pneumacult and Supplemental Figure S3c for BEGM). Cell trajectories and transitions from one cell population to another were deduced from a trajectory inference analysis using Monocle 2, followed by differential expression analysis between consecutive cell states in pseudotime using Seurat. In the BEGM condition, a unique cell trajectory was found (Supplemental Figure S3d), starting with cycling and non-cycling BCs at its beginning, followed by *KRT5*+ and then *KRT5*-supraBCs cells, and with MCCs at its end. Before reaching the secretory-like cell state, the trajectory goes through the 2 “undefined intermediate” cell states. Despite the absence of *SCGB1A1* expression in secretory-like cells (*SCGB1A1*-/*BPIFA1*+/*KRT8*+), these cells were ordered in the pseudotime before MCCs, as expected for canonical SCs (Supplemental Figure S3d-f). A more complex trajectory was observed with the Pneumacult condition, in which Monocle2 detected a bifurcation into 2 distinct branches after the SC stage: the larger branch leading to *FOXJ1*+ MCCs, and the smaller one leading to *MUC5AC*+ GCs (Fig. 1f, g). A closer examination of the pseudotime ordering and differential gene expression (Fig. 1h) revealed that few *MUC5AC*+ cells were found on the MCC branch, after the GC bifurcation and that some *FOXJ1*+ cells retained expression of *MUC5AC*. Altogether, our findings confirm SCs as precursors of both MCCs and GCs. They also suggest that GCs can also act as MCC precursors in airway epithelial regeneration.

### Goblet cells can act as differentiation intermediates for multiciliated cells

We further investigated the possibility that some GCs may correspond in fact to precursor cells for MCCs. A first evidence is coming from our robust clustering analyses, either when they were performed on cells coming from *in vitro* samples or from fresh tissues. The two populations of GCs and SCs displayed very similar gene expression profiles, and were discriminated based on their levels of expression of *MUC5AC* and *MUC5B* which were higher in GCs (Supplemental Figure S1a-c). In the Pneumacult experiment, 24 out of the 54 top genes for GCs were also associated with SCs (Fig. 2a). Among these transcripts was *SCGB1A1*, while expression of *MUC5AC* and *MUC5B* was more robust in GCs (Fig. 2b). When we directly assessed differential gene expression between cells located at the two ends of the GC branch (Fig. 1f, Fig. 2c), we confirmed the high similarity of gene expression existing between SCs and GCs. GCs differed from SCs by higher levels of expression for SC/GC specific genes. This was observed for molecules such as mucins (*MUC1*, *MUC4*, *MUC5B*, *MUC5AC*), secretoglobins (*SCGB1B1* and *SCGB3A1*), PLUNC antimicrobial factors (*BPFA1* and *BPIFB1*) and *SLPI*, the secretory leukocyte protease inhibitor (Fig. 2c). These properties led us to consider GCs as ‘hyperactive’ SCs and led to the prediction that these cells could also function as MCC precursors. This point was tested by quantifying the cellular expression of *MUC5AC* and *FOXJ1*, and by measuring the percentage of doubly labeled cells. Our rationale was that a detection of cells that express at the same time MUC5AC and FOXJ1 would suggest the existence of a transitory state between GCs and MCCs. Fig. 2d, 2g and 2j indeed show that 8.9% of GCs + MCCs express at the same time *MUC5AC* and *FOXJ1*. It also shows the existence of SCs/MCCs expressing both *SCGB1A1* and *FOXJ1*, which correspond to a more conventional type of precursors for MCCs (Fig. 2m). The presence of *MUC5AC*+/*FOXJ1*+ and *SCBG1A1*+/*FOXJ1*+ cells was not restricted to a cell culture model, as these transitionary cells were also detected in fresh biopsies from human bronchi (Fig. 2e, h, k, n) and pig trachea (Fig. 2f, i, l, o). The presence of doubly-labeled cells was confirmed by qPCR in a fully independent dataset, derived from a HAEC culture (Supplemental Figure S5). In that other experiment, we isolated the cells with the C1 technology (Fluidigm) and quantified gene expression by quantitative RT-PCR with a Biomark (Fluidigm). Cells isolated with the C1 were visually inspected, and these experimental settings ensured the absence of cell doublets. We found that 4 cells out of 74 expressed GC specific genes (namely *MUC5AC*, *MUC5B* and *TFF3*), together with MCC specific genes (*FOXJ1*), and more specifically, immature MCC genes (*PLK4*, *MYB* and *CDC20B*) [32] (Supplemental Figure S5a, b). The result was further confirmed after reanalyzing a recent dataset published by others [29] (Supplemental Figure S5c). A further confirmation came from the detection at a protein level of cells that were labeled at the same time by MUC5AC and acetylated Tubulin, a specific protein marker of the cilia (Fig. 2p). A final point came after a survey of our data with “RNA velocity” [33]. Velocity can predict the fate of individual cells at a timescale of hours by distinguishing the expression of spliced and unspliced forms of transcripts. We used Velocity to analyze the behavior of three transcripts: *CEP41*, a specific marker of cells in an early phase of multiciliated differentiation, *SCGB1A1* and *MUC5B*. RNA velocity algorithm calculates a residual value of each gene, which indicates expected upregulation when it is positive and expected downregulation when it is negative. Positive residuals were found for transcripts of *CEP41* in the GC population, predicting an upregulation of this gene’s expression in the next hours. A different picture was observed for the transcripts of *SCGB1A1* and *MUC5B*, in which negative residuals were found in the GC and SC populations, indicating an expected downregulation of the corresponding transcripts in the next hours (Fig. 2q). Altogether, these data indicate that GCs can act as precursors for MCCs in normal *in vitro* and in homeostatic *in vivo* airway regeneration.

**Figure 2:**
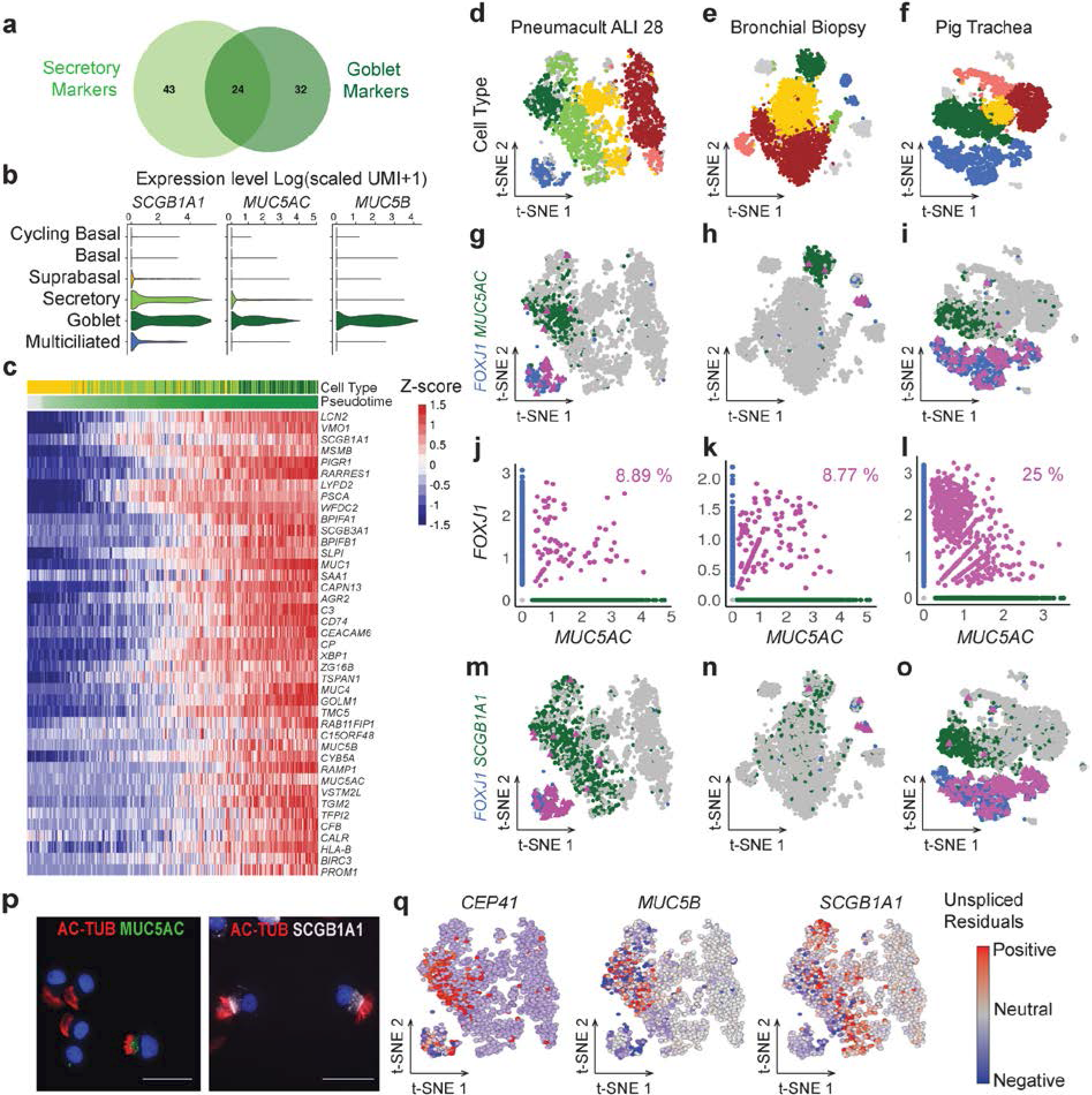
Goblet cells as differentiation intermediates for multiciliated cells. **a** Venn diagram illustrating closeness of the best marker genes for secretory and goblet cells deduced from scRNA-seq of cells differentiated in Pneumacult medium (ALI28). **b** Violin plots of normalized expression for *SCGB1A1*, *MUC5AC*and *MUC5B*, three markers of secretory and goblet cells. **c** Heat map of the most differentially expressed genes between groups of suprabasal, secretory and goblet cells at key point in the pseudotime (before branching, start of the GC branch and end of the GC branch). Cells are ordered by pseudotime. Bars on the top of the heatmap indicate cell type and pseudotime. **d-e-f** t-SNE plots of expression from scRNA-seq of ALI28 (d), bronchial biopsy cells (e), pig tracheal cells (f). Colors indicates cell types as in Figure 1. **g-h-i** Highlights of gene expression for *FOXJ1*+ cells (blue), *MUC5AC*+ cells (green), and *FOXJ1*+*/MUC5AC*+ cells (pink) in the same samples as in d-e-f. **j-k-l** Relationships between normalized expression of *MUC5AC*and *FOXJ1*in the three same samples. **m-n-o** Highlights of gene expressions for *FOXJ1*+ cells (blue), *SCGB1A1*+ cells (green), and *FOXJ1*+/*SCGB1A1*+ cells (pink). **p** Immunodetection of cells co-expressing markers of multiciliated cells (Acetylated Tubulin) and of goblet cells (*MUC5AC*) (o) or of secretory cells (*SCGB1A1*) (p). **q** Representation by a tSNE plot (scRNA-seq of cells differentiated in Pneumacult medium at ALI28) of the Velocity residuals colored according to estimates of the positive (red) and negative (blue) residues for a multiciliated cell marker (*CEP41*), a goblet cell marker (*MUC5B*) and a secretory cell marker (*SCGB1A1*).

### Refining cell clustering identifies 6 additional clusters, including a discrete population of pre-MCC “deuterosomal” cells

To gain further insight in the diversity of cell populations composing the airway epithelium and the transitionary ones occurring during the regeneration, we considered additional clusters that could be derived from our sub-clustering analysis, by accepting less discriminations between them than between the initial 6 previously identified clusters. This deeper analysis defined 12 clusters, rather than the 6 initial ones (Fig. 3a, Supplemental Figure S6a). The non-cycling BC population was split into 2 clusters that we termed BC1 and BC2. The major difference between these 2 clusters was the higher level of expression for genes associated with cell migration: *FN1*, *VIM*, *SPARC* and *TAGLN* in the BC2 cluster. Analysis of enriched canonical pathways with Ingenuity Pathway Analysis showed enrichment for integrin, actin cytoskeleton and Rho GTPase signaling as well as the pathway “regulation of actin-based motility” in BC2 compared to BC1, suggesting an increased migratory activity in BC2 (Supplemental Figure S7). Since BC2 cells were associated with higher pseudotime values than BC1, BC2 likely correspond to the population of most differentiated BCs, just before reaching a suprabasal state. The *supraBC* and SC populations could also be further split into 3 distinct sub-populations, resulting in a total of 3 new populations of supraBC and 3 new populations of SC (Fig. 3a and Supplemental Figure S6a). Each of them displayed its own distinct gene set enrichment (Supplemental Figure S7). Among them, the SC2 subpopulation appeared particularly interesting, since it displayed a strong enrichment score for the feature “immune cell migration, invasion and chemotaxis”, and also a strong positive enrichment for canonical pathways such as “neuroinflammation signaling” and “dendritic cell maturation”. This was explained by the increased gene expression of targets for pro-inflammatory molecules such as TNF, IFNG, NFkB, IL1A/B, IL2, or IL6, as well as decreased gene expression for targets for the anti-inflammatory PPARG pathway (Supplemental Figure S7). This may confer to this subpopulation of SCs a unique relationship with the immune response. The MCC cluster, containing *FOXJ1*+ cells, was further split in 2 discrete clusters: (1) the largest one is positive for mature MCC genes such as *DNAH5*, and corresponds to terminally differentiated MCCs; (2) the second one specifically expresses several molecules that are important for the biosynthesis of hundreds of motile cilia. Among them is *DEUP1*, a hallmark of massive centriole amplification at deuterosomes (Fig. 3b). This led us to coin these cells “deuterosomal” cells. This subpopulation is clearly distinct from mature MCCs in our clustering analysis (Fig. 3b) and expresses also highly specific markers such as *PLK4*, *CCNO*, and *CEP78* (Supplemental Figure S8). We have confirmed the existence of deuterosomal cells in three distinct species with single cell experiments performed on mouse tracheal epithelial cells (MTECs) dissociated at ALI3 (i.e. the time point of maximal centriole amplification in this model), newborn pig trachea and human bronchial biopsy from a healthy adult subject (Fig. 3c). In all samples, even under homeostatic conditions, we noticed the presence of deuterosomal cells that clustered independently of mature MCCs. This deuterosomal cell population was characterized by the expression of gene markers, either uniquely expressed in deuterosomal cells, or also expressed in MCCs and cycling BCs (Fig. 3d). Our analysis shows that the group of deuterosomal cells specifically expressed 149 genes, while sharing 33 and 244 genes with cycling BCs and mature MCCs, respectively (Fig. 3e). Among the 33 genes in common with cycling BCs, we noticed the re-expression of several cell cycle-related genes, which are required for the massive amplification of centrioles that takes place [34, 35]. The most specific genes are displayed in Fig. 3d. This analysis not only confirmed the known expression of *CDK1* in deuterosomal cells [35], as it also highlights the expression in deuterosomal cells of genes coding for centromere proteins (*CENPF*, *CENPU*, *CENPW*), securin (*PTTG1*), a core subunit of the condensing complex (*SMC4*) and cyclin-dependent kinases regulatory subunits (*CKS1B*, *CKS2*). We confirmed the deuterosomal-specific expression of *CDC20B*, the miR-449 host gene. We have recently shown that CDC20B is a key regulator of centriole amplification by deuterosomes [32]. Incidentally, we noticed the existence of a novel and short isoform of this gene, arising from alternative splicing and results in the inclusion of a novel exon after the location of the miR-449 family (Fig. 3b and Supplemental Figure S9a, b). We found that this short CDC20B isoform was also detectable in mouse RNA-seq data (Supplemental Figure S9c). Comparison of transcript abundance in several samples, including the Pneumacult ALI28 and the human bronchial biopsy showed higher levels for short *CDC20B* compared to *CDC20B*(Supplemental Figure S9d). This short *CDC20B* likely corresponds to the major source of miR-449 in deuterosomal cells. A list of novel markers of deuterosomal cells that are specifically expressed in this cell population is provided in Supplemental Table S1. Some of these genes had never been described before in the context of centriole amplification, such as the Yippee-like factor YPEL1 or the Notch pathway related hairy-enhancer-of-split family of transcription factors HES6 (Supplemental Figure S8). Gene set enrichment of the deuterosomal population specific genes (Fig. 3f) shows an enrichment for terms such as cilium assembly, centrosome maturation but also, cell-cycle mechanism-related terms such as resolution of sister chromatid cohesion, regulation of AURKA and PLK1 activity, CDH1 autodegradation. “Mitochondrial membrane part” was also among the enriched terms, suggesting an increase in mitochondria numbers at this stage. This deuterosome-specific signature perfectly delineates the events that are occurring at this MCC differentiation stage and provides an extensive repertoire of specific cell-cycle related genes which are re-expressed at the deuterosomal stage. This deuterosomal population is a consistent population, much larger than recently described rare cell populations such as ionocytes [28, 29], which we also identified (Supplemental Figure S6c).

**Figure 3:**
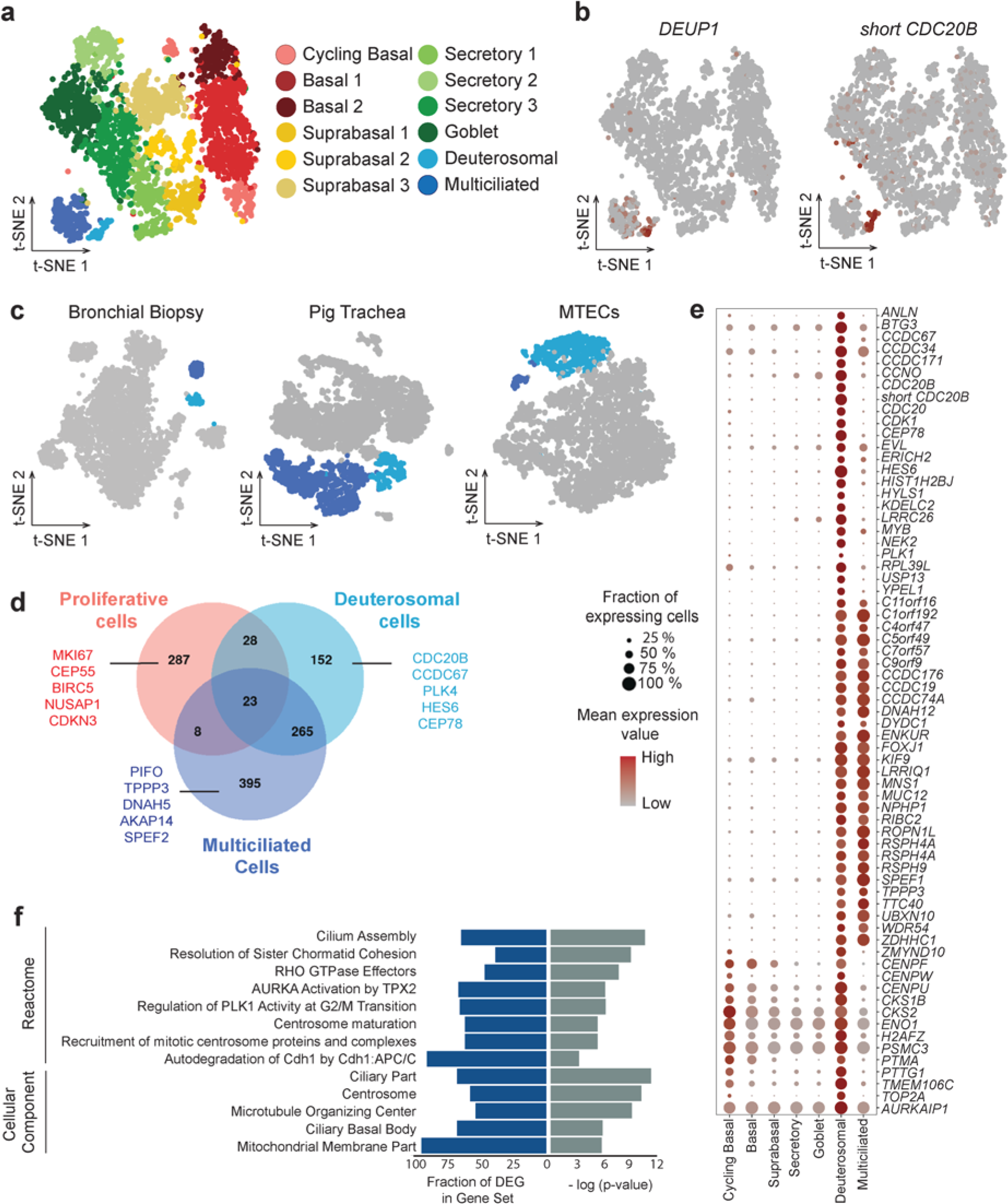
Deuterosomal cells form a discrete MCC intermediate population with a centriole amplification signature. **a** Subcluterization of scRNA-seq from cells differentiated in Pneumacult medium (ALI28) into 12 cell types, deduced from intra-heterogeneity analysis of the 6 initial clusters. **b** Illustration of the specific expression of *DEUP1*and *short CDC20B*in the deuterosomal cell population (low to high expression, grey to red). **c** Identification of the cluster of deuterosomal cells in scRNA-seq data from a biopsy of human bronchi, pig trachea and mouse primary culture (MTEC, ALI3). Deuterosomal (light blue). Multiciliated cluster (dark blue). **d** Venn Diagram showing overlaps existing between top gene markers of deuterosomal cells (light blue) and those of proliferative (pink) or multiciliated cells (dark blue). **e** Dot plot of marker genes for the deuterosomal cell population. Color gradient (grey to red) and dot size indicate for each cluster the mean marker expression and the percentage of cells expressing the marker, respectively. **f** Enriched gene sets in deuterosomal cell marker genes.

### Establishing a keratin switch pattern during airway regeneration

An extremely diverse repertoire of keratins has been associated with different types of epithelial cells. Considering that expression of specific keratins varies with cell type, period of embryonic development, stage of histologic differentiation, cellular growth environment, and disease state, we reasoned that it would be useful to compare our scRNA-seq data with the expression of different keratins. *KRT5* and *KRT14*, are largely used as BC markers in the airway and lung epithelia, and also in other epithelia such as bladder [36], prostate [37] or mammary gland [38]. KRT8 is clearly associated with luminal cell types [6]. Besides, there is no clear repertoire of the other keratins that are associated with airway cell types. A recent study performed on mouse and human models of *in vitro* regeneration identified *KRT4* and *KRT13* in a subpopulation reminiscent of our supraBCs, as it emerges between BCs and SCs [29]. We have established an extensive single-cell repertoire of KRT expression during airway regeneration based on pseudotime ordering in our Pneumacult ALI28 and pig trachea datasets. Our analysis confirmed in both datasets the presence of *KRT5* and *KRT14* in BCs, of *KRT4* and *KRT13* in supraBCs, and the expression of *KRT8* in luminal cell types (SCs, GCs and MCCs) (Fig. 4a and e). We consistently noticed that expression profiles of *KRT13* and *KRT4* did not completely overlap, with *KRT13* detected at earlier pseudotimes than *KRT4*. This was confirmed at a protein level by a quantification by immunostainings of the proportion of KRT5+/KRT13+ and KRT5+/KRT4+ double positive cells (Fig. 4b). Fig. 4c shows that there was more KRT5+/KRT13+ (7.4%) than KRT5+/KRT4+ (4.9%) double positive cells, which is consistent with an earlier expression of KRT13 compared to KRT4. In the pig trachea, we also found a very clear shift, with 16.8% and 11.2% of *KRT5*+/*KRT13*+ and *KRT5*+/*KRT4*+ double positive cells, respectively (Fig. 4d and Supplemental Figure S10). Our results show that *KRT4* and *KRT13* are not strictly expressed at the same time during airway regeneration and their expression delineate subtle differences in cell subpopulations. Additional keratins such as *KRT16* and *KRT23* displayed a specific supraBC expression (Fig. 4e). We also identified additional keratins that were more specifically associated to more differentiated cell types: *KRT7* and *KRT18* were strongly enriched in SCs, but their expression completely dropped in MCCs, while *KRT8* was still expressed (Fig. 4e). Expression patterns for these cell type specific keratins were confirmed in the pig trachea (Supplemental Figure S10c). The keratin switch pattern is indeed sufficiently specific to allow a reconstruction of the cell trajectories during airway regeneration.

**Figure 4:**
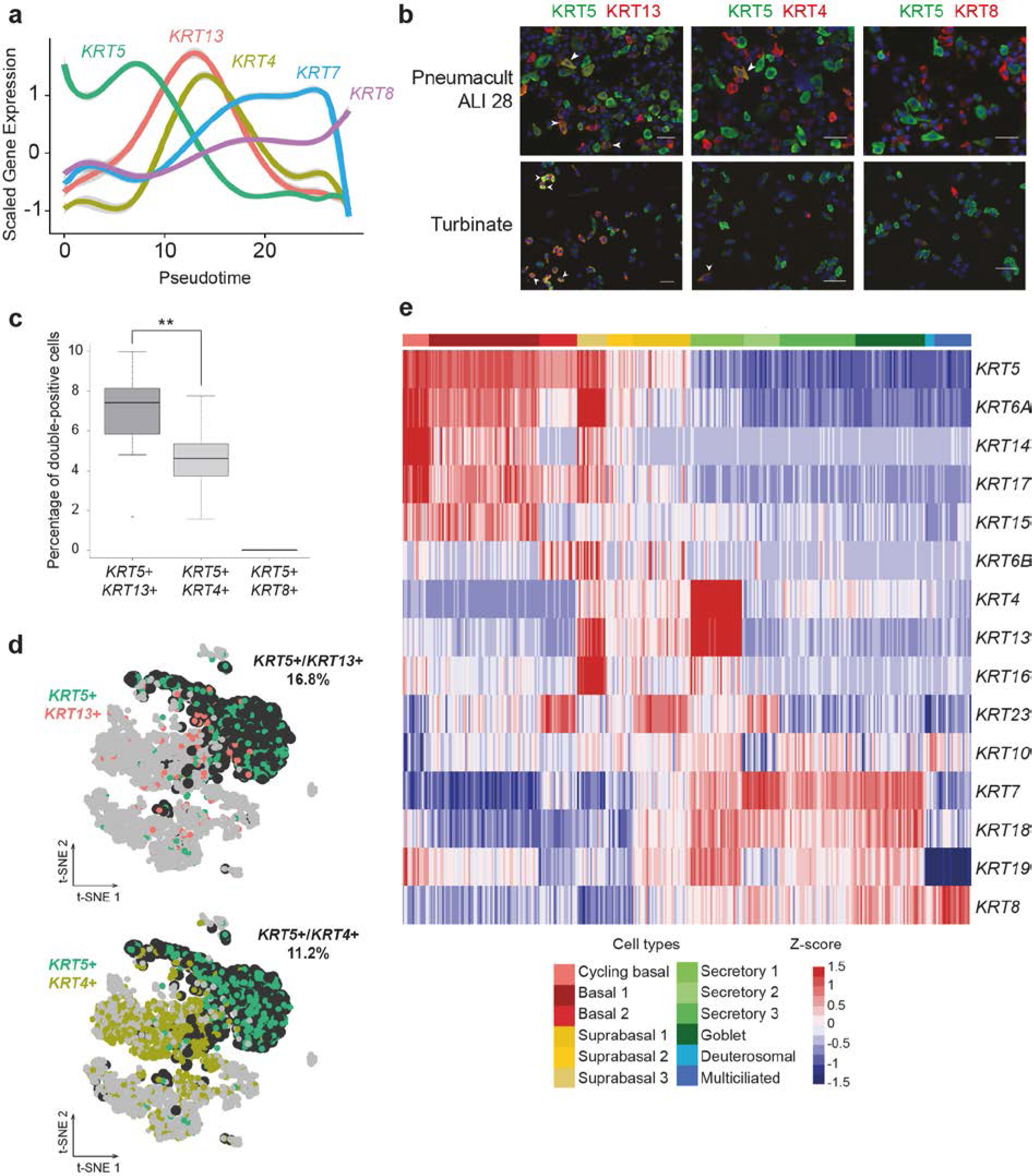
Keratin signatures switch during the differentiation process. **a** Plot of normalized gene expression of keratins according to pseudotime from scRNA-seq of cells differentiated in Pneumacult medium (ALI28). **b** Double immunofluorescence staining for KRT5 and KRT13, KRT4 or KRT8. White arrowheads indicate doubly labelled cells (KRT5+/KRT13+, KRT5+/KRT4+, KRT5+/KRT8+). Nuclei are shown in blue (DAPI). **c** Quantification of double-positive cells from b. **: pvalue<0.01 (Wilcoxon test). **d t**SNEs of scRNA-seq data from pig tracheal epithelial cells. KRT5+ cells are shown in green, KRT13+ are shown in red, KRT4+ are shown in light green, double-positive cells are shown in black. The indicated percentage corresponds to double-positive cells. **e** Heatmap for scRNAseq data from Pneumacult ALI28 showing gene expression for keratins.

### Establishing a combinatorial repertoire of signaling pathways during airway regeneration

We have finally analyzed the cell specificity of expression of important signaling pathways, to catch mutual influences between distinct cells that could play a role in airway regeneration. Our investigation was focused on the Notch, BMP/TGFβ and Wnt pathways. For each different component, we classified them as ligands, receptors, or targets. The expression profiles are shown as heatmaps, cells being sorted by their subgroups, then by ascending pseudotime value (Fig. 5a-c).

**Figure 5:**
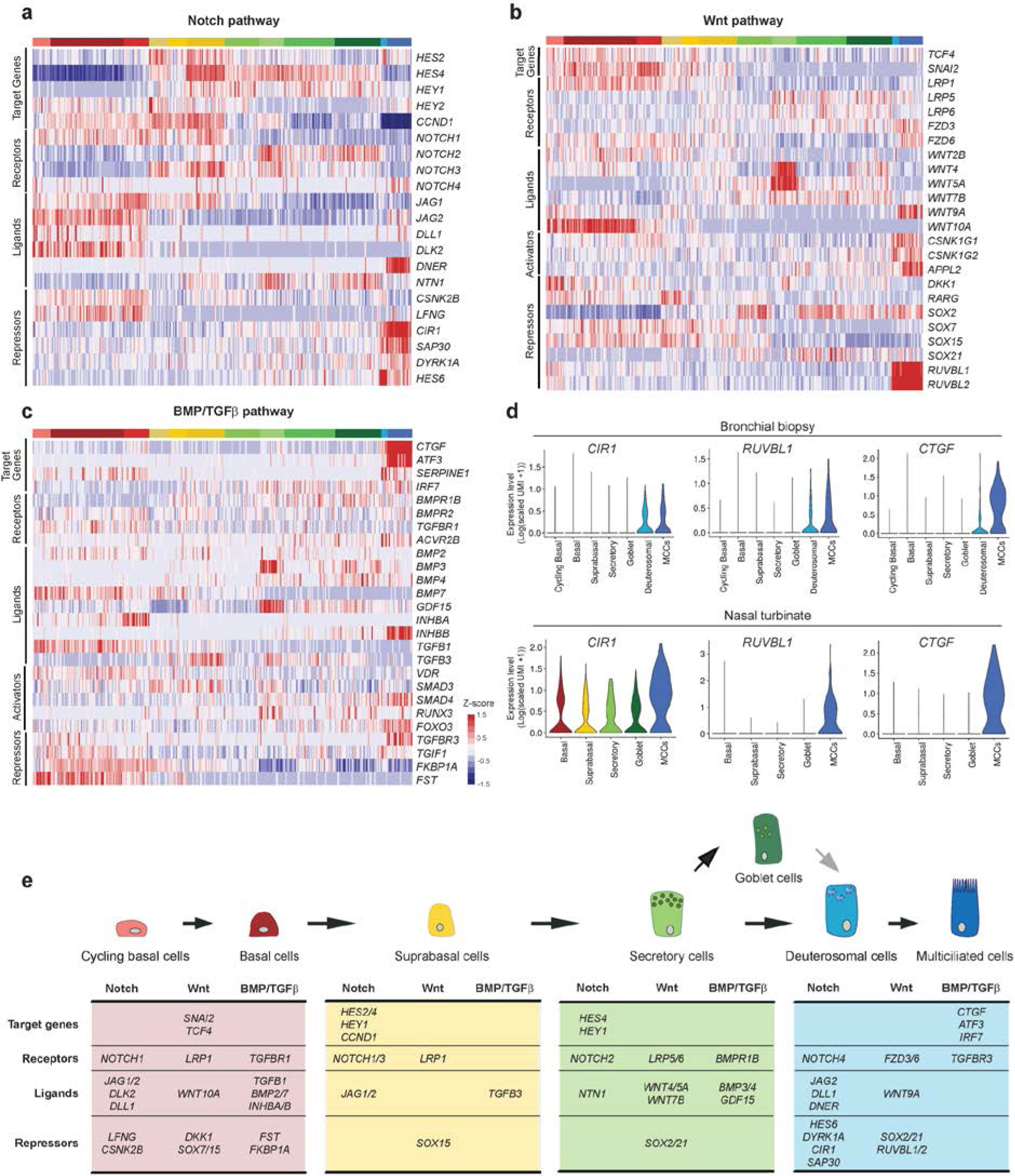
Single-cell expression of signaling pathways components during airway regeneration. **a** Heatmap of the genes related to the Notch pathway] with cells ordered by cluster. **b** Heatmap of the genes related to the Wnt pathway with cells ordered by cluster. **c** Heatmap of the genes related to the BMP/TGFβ pathway with cells ordered first by cluster. **d** Violin plots for selected genes in the bronchial biopsy and nasal turbinate samples. **e** Summary of the major partners involved in specific cell types for the three pathways. Cell type colors are as in Fig. 3 and 4.

#### Notch pathway (Fig. 5a).

BCs express the ligands *DLL1*, *JAG1* and *JAG2*, as well as the receptor *NOTCH1*, as expected [6, 29]. In this population, no target gene expression is detected, suggesting that the pathway is inactive. Interestingly, BCs also express *LFNG*, which has been described to inhibit JAG1 signaling via NOTCH1 [39]. SupraBCs cells express *NOTCH1*, *JAG1* and *JAG2* but, as opposed to BCs, they show clear activation of the Notch pathway by expression of the target genes *HEY1*, *HES2* and *HES4*. *NOTCH3* expression is turned on and is specific to this population. In SCs/GCs, *NOTCH2* is the major receptor to be detected and signal activation remains as evidenced by the expression of *HEY1* and *HES4*. SCs/GCs also express the non-canonical Notch ligand *NTN1*. In Deuterosomal cells/MCCs, a clear shift is observed. *NOTCH2*, *NOTCH3*, *HEY1* and *HES4* are turned down, and *NOTCH4* is specifically expressed. As previously described, *JAG2* [29], which is present in BCs, then absent in supraBCs and SCs/BCs, is re-expressed in the MCC compartment. We have found the same behavior for *DLL1* and the non-canonical ligand *DNER1*. Thus, MCC express some Notch ligands. Strikingly, a major inhibitory signature dominates in MCCs, with the expression of *CIR1* and *SAP30*, which are transcriptional corepressors, as well as of *DYRK1A*, an inhibitor of NICD. *HES6*, which expression is not regulated by Notch signaling but which has been identified as a Notch pathway inhibitor through its binding to HES1 [40], displays an expression pattern that is highly enriched in deuterosomal cells (Fig. 5a and Fig. 3d).

#### Wnt pathway (Fig. 5b).

The Wnt target genes *SNAI2* and *TCF4*, indicators of an active pathway are mainly enriched in the BC population, especially in BC2 for SNAI2. In the BC population, *WNT10A* and *LRP1* are strongly enriched, and several *SOX* family members (*SOX2* and *SOX21*) are underrepresented, especially in the cycling BCs, suggesting an activation of the pathway in this compartment. In the MCC population, the situation is more complex. Despite a slight expression of *TCF4* together with positive regulators of the pathway such as *WNT9A*, *FZD6*, *APPL2*, *CSNK1G1* (a casein kinase component that can act as an activator or inhibitor of the pathway [41]), no *SNAI2* expression is detected and known repressors of the Wnt pathway are also overrepresented. Indeed, MCCs express significant levels of the transcriptional repressors *SOX2*, *SOX21* and display strong enrichments for the Reptin components *RUVBL1* and *RUVBL2* (Fig. 5b).

#### BMP/TGF*β* pathway (Fig. 5c).

BMP ligands, such as *BMP2*, and *BMP7*, are both enriched in the BC population, while *BMP3* and *BMP4* are both enriched in the SC/GC populations. A different picture was found for BMP receptors, for which we did not found specific cell population expression. The most striking observation is the specific expression in BCs of BMP inhibitors such as the inhibitory ligand FST and the intracellular inhibitor FKBP1A (also known as FKBP12). As BMP signaling is considered as a brake for proliferation, inhibition in the BC compartment is consistent with the maintenance of a proliferative potential of this progenitor population. Regarding the TGFβ pathway, a clear signal of activation is detected in the deuterosomal/MCC population, with specific expression of the target genes *SERPINE1* (PAI-1), *CTGF*, *ATF3*, *TGFBR3* and *IRF7*, consistent with the previous finding that TFGβ pathway regulates motile cilia length by affecting the transition zone of the cilium [42]. We did not detect TGFβ ligands in the MCC population but rather found them expressed in BCs (TGFB1) and supraBCs (TGFB3).

We have confirmed the main distribution of the three pathway components sample differentiated with the BEGM medium (Supplemental Figure S11) and in 2 fresh tissue samples (human bronchial biopsy and nasal turbinate) for which a selection of genes is shown in Fig. 5d. Collectively, our data provide for the first time a detailed repertoire of signaling pathways at work during airway regeneration, with receptors and ligands specifically expressed at each cell stage.

## Discussion

In this study, we have established a comprehensive single-cell atlas throughout the entire time course of human airway regeneration *in vitro*. We carefully chose representative time points in the process and we quantified the proportion and identity of each cell population at different time points after the establishment of the air liquid interface. We provide here for the first time a comparison of the most widely used culture media in 3D culture of airway epithelial cells: BEGM, the most established commercial medium with which the majority of studies have been performed, and a more recent commercial medium, Pneumacult. In the BEGM medium, we have performed analyses at earlier time points, i.e. ALI2 and ALI4. These time points allowed us to measure the extent of cell proliferation during *in vitro* regeneration. Cycling basal cells account for approximatively 40% of total cells at ALI2 and ALI4, and this number drops to 5% at ALI7. These early time points also showed that suprabasal cells appeared early in these conditions, as they were already detected at ALI4. With the BEGM medium, unlike with the Pneumacult medium, we never detected any goblet (*MUC5AC*+) nor “canonical” secretory (secretoglobin+) cell, even after long periods of time, and on several dozens of cultures coming from distinct donors (Supplemental Figure S1, S3, and personal observations). However, we found a cell population that we have termed “secretory-like”. “Secretory-like” cells express a gene pattern very similar to that of canonical secretory cells, and they can differentiate into MCCs. Interestingly, goblet cells were detected in BEGM medium after an IL-13 treatment [41, and our personnal observations]. Future work should investigate whether secretory-like cells first evolve into canonical secretory cells and then goblet cells upon IL-13 treatment.

In the Pneumacult medium, but also in freshly dissociated human bronchial biopsy and pig trachea, we have detected cells expressing both *MUC5AC* and *FOXJ1*. We believe this finding is in line with our lineage inference and RNA velocity data showing that goblet cell can be precursors for MCCs. Other groups had previously detected cells expressing both these markers, in the context of GC hyper/metaplasia induced by Sendai virus infection or after IL-13 treatment [44-46]. These findings led them claim that MCC can transdifferentiate into GCs. However, none of their data clearly showed a difference in the number of these double-positive cells between control and treated conditions. Our present finding probably provide a different answer to this conundrum, saying that these double-positive cells do exist in the absence of IL-13 stimulation. We noticed that their expression profiles posit them more naturally as precursors of MCCs than as trans-differentiated multiciliated cells. Turner and colleagues [45] proposed that MCCs could trans-differentiate into GCs after performing *in vitro* lentiviral transduction of HAECs with a vector containing a Cre recombinase under the control of the *FOXJ1* promoter. These findings were not confirmed by Rajagopal’s group who showed no GC arising from MCC in a context of OVA-induced mucous metaplasia in mouse airways, using *in vivo* lineage tracing with *Foxj1*-cre mice [47].

Our datasets are not limited to HAECs, but also provide useful information about mouse, human, and pig tissues. They correspond to *in vitro* MTECs, as well as cells freshly isolated from human nasal brushing, turbinate, bronchial biopsy and finally pig trachea. We have used this variety of samples to confirm our findings with HAECs, and to invalidate the possibility of culture artefacts. Our study was not focused on rare cell types such as pulmonary neuroendocrine, brush cells or ionocytes, which have been recently described elsewhere. We have indeed detected cells, which displayed the gene signature of ionocytes, characterized by expression of *CFTR* and other ion transporters, as well as specific transcription factors such as *ASCL3* or *FOXI1* [28, 29]. Our investigation was rather focused on the diversity of the main cell types that compose the epithelium. We have identified 3 subtypes of basal cells among which cycling basal cells, and a group of basal cells expressing significantly higher levels of genes involved in extracellular matrix connection and actin-based motility. As these cells are the latest basal cells in our pseudotime, they likely represent an important transient migratory state on the way to the suprabasal state.

An additional peculiarity found within the secretory compartment comes from one of our three secretory subpopulations which displayed an immune-related gene signature. So far, diversity within the secretory compartment was established after expression of different members of the secretoblogin family [48] or via activation level of the Notch pathway [49]. We propose that diversity within the secretory cell compartment should also include specialized functions related to the interaction between the epithelium and immune cells. Additional experiments including protein labeling on fresh tissue sections have now to be performed in order to confirm this diversity and identify the spatial distribution of these subpopulations.

Our study has also provided a first extensive gene signature of the deuterosomal population, which play a key role during differentiation of multicilated cells. In line with what has been shown recently by our group and others [32, 35], cell-cycle related genes become re-expressed in this population on non-cycling cells. We have confirmed the very specific expression of *CDC20B*, a key player of centriole amplification [32], and have identified, both in human and mouse, a novel isoform of this transcript which displays higher expression than the annotated long isoform. As the pre-mRNA corresponding to this short isoform comprises the *miR-449-*encoding intron, we suggest that this isoform should indeed be the major source of *miR-449* in deuterosomal cells. The alternative splicing that is responsible for this alternative isoform might represent an optimization of gene expression regulation to efficiently increase miR-449 levels.

Besides the identification of each cell type, we have also characterized the distribution of some important signaling pathway components. First, we have focused on the Notch pathway as it is already known as a major regulator of the mucociliary differentiation. We have confirmed the distribution of ligands and receptors described by others [19, 20, 29, 50]. We have also confirmed the absence of Notch activation in BCs and MCCs, with *HES4* being the most representative target gene in our model. BCs express *NOTCH1* and NOTCH ligands. However, no clear Notch pathway activation can be detected within this cell population even in an uneven manner as it could be expected from Notch lateral inhibition. This could be solely the result of weak *NOTCH1* expression but could also be a consequence of specific expression of Notch inhibitors such as the ligand *LFNG* or Casein kinase II subunit beta (*CSNK2B*) [51, 52]. At the other end of mucociliary differentiation, inhibition of the Notch pathway has been widely documented in MCCs: failure to inactivate Notch results in failure of MCC signaling differentiation. Here we suggest a novel mechanism for Notch pathway inactivation by showing specific expression of several Notch transcriptional inhibitors at the deuterosomal stage. These include *HES6*, an inhibitory HES acting through HES1 binding [40, 53], *DYRK1A*, an inhibitor of Notch Intracellular Domain transcriptional activity [54], as well as *CIR1* and *SAP30* which are transcriptional repressors of Notch/CSL transcriptional complex [55]. As opposed to MCCs and BCs, SCs must undergo clear Notch activation to maintain their cell identity, and differentiate into GCs [18-20]. However, the onset of this signal’s activation has not been widely studied. Mori and colleagues have described NOTCH3 expression in TP63-negative cells in a parabasal position of the epithelium, which likely correspond to the cells we, and others, have termed supraBCs [50]. We have confirmed that *NOTCH3* RNA is absent from BCs and becomes upregulated in supraBCs. We went further by showing *HES4* becomes expressed at this cell stage, confirming that Notch pathway activation starts at the supraBC stage. Thus, we emphasize here the importance of this intermediate cell population, though it has not been well characterized so far, for establishing Notch activation and subsequent differentiation. Contrary to the Notch pathway, the Wnt/β-catenin pathway has not been extensively studied in the context of airway epithelium differentiation. Crosstalk between these 2 pathways has at least been suggested by other studies performed in other systems. For instance, in the hair follicle precortex, β-catenin stimulates Notch signaling by inducing *Jag1* transcription [56]. In the context of airway epithelium, a very recent study demonstrated that β-catenin signaling is required for the early stages of mucus and ciliated cell differentiation, which they defined as “specification”, but was detrimental to the later “commitment” stages [57]. Wnt also seems to be related to epithelial remodeling upon inflammatory situations: Ordovas-Montanes and colleagues have recently shown that in nasal polyps, which are characterized by an inflammatory state favoring GCs at the expense of MCCs, there is an altered balance between Wnt and Notch signaling, in favor of Wnt signaling [30]. *WNT5A* has been associated with remodeling of airway smooth muscle cells in the context of airway hyperresponsiveness [58]. *WNT4* has been shown to be upregulated in the epithelium of patients suffering from chronic obstructive pulmonary disease and upregulates *IL8* and *CXCL8* gene expression in HAECs [59]. Interestingly, we found both *WNT5A* and *WNT4* specifically expressed by the subpopulation of SCs that may be involved in immune response. This finding again suggests a role for this SC population in the inflammation-induced airway remodeling.

Based on expression of the target genes *TCF4* and *SNAI2*, in our study, activation of the Wnt pathway appears to be confined to the BC population. This population also expresses strongly and specifically the ligand *WNT10A*, suggesting an autocrine regulatory loop. WNT10A has been shown to be specific of BC in other epithelia, such as the mammary epithelium [60]. In fallopian organoids, Wnt has been shown to be essential for stemness [61] and for self-renewal, but not proliferation, in basal-like breast cancer cells [62]. Thus, autocrine WNT10A signaling may regulate self-renewal in the BC compartment of the airway epithelium. In MCCs, we have observed a specific expression of both members from the Reptin family which are ATP-dependent DNA helicases that have been identified as Wnt signaling repressors [63, 64]. As Reynolds and colleagues recently showed that β-catenin was necessary for mucus and ciliated cell specification [57], additional investigations should certainly be carried out to characterize precisely the role of Wnt/β-catenin during airway epithelial regeneration.

## Conclusions

Our lineage inference study during the airway epithelium regeneration has provided novel insights in differentiation dynamics by positioning goblet cells as possible precursors for multiciliated cells. Thus, in the airway epithelium, unlike in most tissues, cells carrying specialized function, i.e. secretory and goblet cells, can constitute differentiation intermediates for other specialized cells, the multiciliated cells. We have also identified subpopulations within the basal, suprabasal, secretory and multiciliated cell compartments. In particular, we propose that if all secretory cells produce anti-microbial peptides, one subset of secretory cells may be more specifically engaged in immune-cell signaling. Our dataset also provides extensive characterization of the deuterosomal cell population. In addition, we have established an exhaustive repertoire of keratin expression and showed that observation of the “keratin switch” during differentiation could be self-sufficient to establish the identity of the different cell populations. Finally, we have improved signaling pathway characterization by detecting putative Notch repressors that might participate in Notch signal shutdown at the deuterosomal stage, and reporting Wnt pathway activity within the basal cell compartment.

## Methods

### Human Airway Epithelial Cells (HAECs) culture

HAECs cultures were derived from nasal mucosa of inferior turbinates. After excision, nasal inferior turbinates were immediately immersed in Ca2+/Mg2+-free HBSS supplemented with 25 mM HEPES, 200 U/mL penicillin, 200 µg/mL streptomycin, 50 µg/mL gentamicin sulfate, and 2.5 µg/ml amphotericin B (all reagents from Gibco). After repeated washes with cold supplemented HBSS, tissues were digested with 0.1% Protease XIV from *Streptomyces Griseus* (Sigma-Aldrich) overnight at 4°C. After incubation, fetal calf serum (FCS) was added to a final concentration of 10%, and nasal epithelial cells were detached from the stroma by gentle agitation. Cell suspensions were further dissociated by trituration through a 21G-needle and then centrifuged at 150g for 5 min. The pellet was resuspended in supplemented HBSS containing 10% FCS and centrifuged again. The second cell pellet was then suspended in Dulbecco’s Modified Eagle’s Medium (DMEM, Gibco) containing 10% FCS and cells were plated (20 000 cells per cm^2^) on 75 cm^2^-flasks coated with rat-tail collagen I (Sigma-Aldrich). Cells were incubated in a humidified atmosphere of 5% CO2 at 37°C. Culture medium was replaced with Bronchial Epithelium Basal Medium (BEBM, Lonza) supplemented with BEGM SingleQuot^TM^ Kit Supplements (Lonza) on the day after and was then changed every other day. After 4 to 5 days of culture, after reaching about 70% confluence, cells were detached with trypsin-EDTA 0.05% (Gibco) for 5 min and seeded on Transwell^®^ permeable supports (6.5 mm diameter; 0.4 µm pore size; Corning), in BEGM medium, with a density of 30 000 cells per Transwell^®^. Once the cells have reached confluence (typically after 5 days), they were induced to differentiate at the air-liquid interface by removing medium at the apical side of the Transwell^®^, and by replacing medium at the basal side with either DMEM:BEBM (1:1) supplemented with BEGM SingleQuot^TM^ Kit Supplements or with Pneumacult-ALI (StemCell Technologies) as indicated in the figure legends. Culture medium was changed every other day.

### Mouse tracheal epithelial cells (MTECs)

MTECs cell cultures were established from the tracheas of 12 weeks-old mice. After dissection, tracheas were placed in cold DMEM:F-12 medium (1:1) supplemented with 15 mM HEPES, 100 U/mL penicillin, 100 µg/mL streptomycin, 50 µg/mL gentamicin sulfate, and 2.5 µg/ml amphotericin B. Each trachea was processed under a binocular microscope to remove as much conjunctive tissue as possible with small forceps and was opened longitudinally with small dissecting scissors. Tracheas were then placed in supplemented DMEM:F-12 containing 0.15% protease XIV from Streptomyces Griseus. After overnight incubation at 4°C, FCS was added to a final concentration of 10%, and tracheal epithelial cells were detached by gentle agitation. Cells were centrifuged at 400g for 10 min and resuspended in supplemented DMEM:F-12 containing 10% FCS. Cells were plated on regular cell culture plates and maintained in a humidified atmosphere of 5% CO2 at 37°C for 4 hours to allow attachment of putative contaminating fibroblast. Medium containing cells in suspension was further centrifuged at 400g for 5 min and cells were resuspended in supplemented DMEM:F-12 containing BEGM Singlequot^TM^ kit supplements and 5% FCS. Cells were plated on rat tail collagen I-coated Transwell^®^. Typically, 5 tracheas resulted in Transwells^®^. Medium was changed every other day. Air-liquid interface culture was conducted once transepithelial electrical resistance had reached a minimum of 1000 ohm/cm2 (measured with EVOM2, World Precision Instruments). Air-liquid interface culture was obtained by removing medium at the apical side of the Transwell^®^, and by replacing medium at the basal side with Pneumacult-ALI medium (StemCell Technologies).

### HAEC and MTEC dissociation for single-cell RNA-seq

Single-cell analysis was performed at the indicated days of culture at the air-liquid interface. To obtain a single-cell suspension, cells were incubated with 0.1% protease type XIV from *Streptomyces griseus*(Sigma-Aldrich) in supplemented HBSS for 4 hours at 4°C degrees. Cells were gently detached from Transwells^^®^^ by pipetting and then transferred to a microtube. 50 units of DNase I (EN0523 Thermo Fisher Scientific) per 250 µL were directly added and cells were further incubated at room temperature for 10 min. Cells were centrifuged (150g for 5 min) and resuspended in 500 µL supplemented HBSS containing 10% FCS, centrifuged again (150g for 5 min) and resuspended in 500 µL HBSS before being mechanically dissociated through a 26G syringe (4 times). Finally, cell suspensions were filtered through a 40 µm porosity Flowmi™ Cell Strainer (Bel-Art), centrifuged (150 g for 5 min) and resuspended in 500 µL of cold HBSS. Cell concentration measurements were performed with Scepter™ 2.0 Cell Counter (Millipore) and Countess™ automated cell counter (Thermo Fisher Scientific). Cell viability was checked with Countess™ automated cell counter (Thermo Fisher Scientific). All steps except the DNAse I incubation were performed on ice. For the cell capture by the 10X genomics device, the cell concentration was adjusted to 300 cells/µl in HBSS aiming to capture 1500 cells for HAECs and 5000 cells for MTECs.

### Turbinate epithelial cell dissociation

To obtain a single-cell suspension from turbinates, the whole turbinate was incubated with 0.1% protease type XIV from *Streptomyces griseus (*Sigma-Aldrich) in supplemented HBSS at 4°C degrees overnight. Epithelial cells were gently detached from the turbinate by washing with HBSS pipetting up and down and then transferred to a 50 ml Falcon tube. Cells were centrifuged (150g for 5 min at 4°C), after removing of the supernatant the cells were resuspended in 1 ml of HBSS, 50 units of DNase I (EN0523 Thermo Fisher Scientific) per 250 µL were directly added and cells were further incubated at room temperature for 10 min. Cells were centrifuged (150g for 5 min at 4°C) and resuspended in 1 ml supplemented HBSS containing 10% FCS, centrifuged again (150g for 5 min at 4°C) and resuspended in 500 µL HBSS before being mechanically dissociated through a 26G syringe (4 times). Finally, cell suspensions were filtered 40 µm porosity Flowmi™ Cell Strainer (Bel-Art), centrifuged (150 g for 5 min) and resuspended in 500 µL of cold HBSS. Cell concentration measurements were performed with Scepter™ 2.0 Cell Counter (Millipore) and Countess™ automated cell counter (Thermo Fisher Scientific). Cell viability was checked with Countess™ automated cell counter (Thermo Fisher Scientific). All steps except the DNAse I incubation were performed on ice. For the cell capture by the 10X genomics device, the cell concentration was adjusted to 500 cells/µl in HBSS aiming to capture 5000 cells.

### Anesthetic procedure

Intranasal anesthesia is performed with topical application (gauze) of 5% lidocaine (anesthetic) plus naphazoline (vasoconstrictor) solution (0.2 mg/ml). Laryngeal and endobronchial anesthesia is performed with topical application of 2% lidocaine through the working channel of a 4.9 mm outer diameter bronchoscope.

### Nasal brushing

Brushing was performed with a 2 mm cytology brush (Medi-Globe) in the inferior turbinate zone.

### Bronchial biopsy

Bronchial biopsy was performed at the spur between the left upper lobe and the left lower lobe with a 1.8mm-diameter Flexibite biopsy forceps (Medi-Globe) passed through the working channel of the bronchoscope (WCB).

### Dissociation of nasal brushing

The brush was soaked in a 5 mL Eppendorf containing 1 mL of dissociation buffer which was composed of HypoThermosol^®^ (BioLife Solutions) 10 mg/mL protease from *Bacillus Licheniformis* (Sigma-Aldrich, reference P5380) and 0.5 mM EDTA. The tube was shaken vigorously and centrifuged for 2 min at 150 g. The brush was removed, cells pipetted up and down 5 times and then incubated cells on ice for 30 min, with gentle trituration with 21G needles 5 times every 5 min. Protease was inactivated by adding 200 μL of HBSS/2% BSA. Cells were centrifuged (400g for 5 min at 4°C). Supernatant was discarded leaving 10 μL of residual liquid on the pellet. Cells were resuspended in 500 μL of wash buffer (HBSS/0.05% BSA) and 2.250 mL of Ammonium Chloride 0.8% was added to perform red blood cell lysis. After a 10 min incubation, 2 mL of wash buffer were added, and cells were centrifuged (400g for 5 min at 4°C). Supernatant was discarded leaving 10 μL of residual liquid on the pellet, cells were resuspended in 1000 μL of wash buffer centrifuged (400g for 5 min at 4°C). Supernatant was discarded leaving 10 μL of residual liquid on the pellet, cells were resuspended in 1000 μL of wash buffer and passed through 40 µm porosity Flowmi™ Cell Strainer (Bel-Art), then centrifuged (400g for 5 min at 4°C). Supernatant was discarded leaving 10 μL of residual liquid on the pellet, cells were resuspended in 100 μL of wash buffer. Cell counts and viability were performed with Countess™ automated cell counter (Thermo Fisher Scientific). For the cell capture by the 10X genomics device, the cell concentration was adjusted to 500 cells/µl in HBSS aiming to capture 5000 cells. All steps were performed on ice.

### Dissociation of bronchial biopsy

The biopsy was soaked in 1 mL dissociation buffer which was composed of DPBS, 10 mg/mL protease from *Bacillus Licheniformis* (Sigma-Aldrich, reference P5380) and 0.5 mM EDTA. After 1 h, the biopsy was finely minced with a scalpel, and returned to dissociation buffer. From this point, the dissociation procedure is the same as the one described in the “dissociation of nasal brushing” section, with an incubation time increased to 1h, and omitting the red blood cell lysis procedure. For the cell capture by the 10X genomics device, the cell concentration was adjusted to 300 cells/µl in HBSS aiming to capture 5000 cells. All steps were performed on ice.

### Pig tracheal epithelial cell dissociation

To obtain a single-cell suspension from pig trachea, whole clean tracheas were incubated with 0.1% protease type XIV from *Streptomyces griseus* (Sigma-Aldrich) in supplemented HBSS at 4°C degrees overnight. Epithelial cells were gently detached from the turbinate by washing with HBSS pipetting up and down and then transferred to a 50 ml Falcon tube. Cells were centrifuged (150g for 5 min at 4°C), after removing of the supernatant the cells were resuspended in 1 ml of HBSS, 50 units of DNase I (EN0523 Thermo Fisher Scientific) per 250 µL were directly added and cells were further incubated at room temperature for 10 min. Cells were centrifuged (150g for 5 min at 4°C) and resuspended in 1 ml supplemented HBSS containing 10% FCS, centrifuged again (150g for 5 min at 4°C) and resuspended in 500 µL HBSS before being mechanically dissociated through a 26G syringe (4 times). Finally, cell suspensions were filtered through 40 µm porosity Flowmi™ Cell Strainer (Bel-Art), centrifuged (150 g for 5 min) and resuspended in 500 µL of cold HBSS. Cell concentration measurements were performed with Scepter™ 2.0 Cell Counter (Millipore) and Countess™ automated cell counter (Thermo Fisher Scientific). Cell viability was checked with Countess™ automated cell counter (Thermo Fisher Scientific). All steps except the DNAse I incubation were performed on ice. For the cell capture by the 10X genomics device, the cell concentration was adjusted to 500 cells/µl in HBSS aiming to capture 5000 cells.

### Single-cell RNA-seq

We followed the manufacturer’s protocol (Chromium™ Single Cell 3′ Reagent Kit, v2 Chemistry) to obtain single cell 3’ libraries for Illumina sequencing. Libraries were sequenced with a NextSeq 500/550 High Output v2 kit (75 cycles) that allows up to 91 cycles of paired-end sequencing: Read 1 had a length of 26 bases that included the cell barcode and the UMI; Read 2 had a length of 57 bases that contained the cDNA insert; Index reads for sample index of 8 bases. Cell Ranger Single-Cell Software Suite v1.3 was used to perform sample demultiplexing, barcode processing and single-cell 3’ gene counting using standards default parameters and human build hg19, pig build sus scrofa 11.1 and mouse build mm10.

### Single-cell quantitative PCR

HAECs were dissociated as described above, then single cells were separated with a C1^TM^ Single-cell AutoPrep system (Fluidigm), followed by quantitative PCR on the Biomark system (Fluidigm) using SsoFast^TM^ evaGreen^®^ Supermix (Biorad) and the primers described in Supplementary Table S2.

### RNA-seq on dissociated and non-dissociated HAECs

Two Transwells^®^ from fully differentiated HAECs from 2 distinct donors were each dissociated as described above. After the final resuspension, cells were centrifuged and resuspended in 800 µL Qiazol (Qiagen). Non-dissociated cells from 2 Transwells^®^ were also lyzed in 800 µL Qiazol. RNAs were extracted with the miRNeasy mini kit (Qiagen) according to manufacturer’s instructions. Two micrograms from each RNA was used in RNAseq library construction with the Truseq^®^ stranded total RNA kit (Illumina). Sequencing was performed with a NextSeq 500/550 High Output v2 kit (75 cycles). Reads were aligned against hg19 human build using STAR aligner. Low expressed genes were filtered out, then paired differential analysis was performed with DESeq2 comparing dissociated vs non-dissociated samples from cultures generated from 2 different donors. P-values were adjusted for multiple testing using the false discovery rate (FDR). Top differentially expressed genes were selected using the following cutoffs: FDR < 0.001 and an absolute log2FC > 1.5

### Cytospins

Fully differentiated HAECs were dissociated by incubation with 0.1% protease type XIV from *Streptomyces griseus* (Sigma-Aldrich) in HBSS (Hanks’ balanced salts) overnight at 4°C. Cells were gently detached from the Transwells^®^ by pipetting and then transferred to a microtube. Cells were then cytocentrifuged at 72 g for 10 min onto SuperFrost^TM^ Plus slides using a Shandon Cytospin^TM^ 4 cytocentrifuge.

### Tissue processing and OCT tissue embedding (see if we maintain figures containing this part)

Nasal turbinates were fixed in PFA 4% at 4°C overnight then, for cryoprotection, tissues were soaked in a 15% sucrose solution until tissue sinks, then soaked in a 30% solution before tissue sinks. Tissue was embedded in “optimal cutting temperature” OCT medium (Thermo Fisher Scientific) at room temperature and then submerged in Isopentane previously tempered at −80°C. Cutting of frozen tissues was performed with a cryostat Leica CM3050 S.

### Immunostaining

Cytospin^TM^ slides were fixed for 10 min in 4% paraformaldehyde at room temperature and then permeabilized with 0.5% Triton X-100 in PBS for 10 min. Cells were blocked with 3% BSA in PBS for 30 min. The incubation with primary antibodies was carried out at 4°C overnight.

Primary antibodies: mouse monoclonal KRT4 (1:50, Santa Cruz Biotechnology sc-52321), mouse monoclonal KRT8 (1:50, Santa Cruz Biotechnology, sc-58737), mouse monoclonal KRT13 (1:200, Sigma-Aldrich clone KS-1A3), rabbit KRT5 (1:2000, Biolegend BLE905501), Rabbit CC10 (SCGB1A1) (1:500, Millipore 07-623), mouse monoclonal Acetylated Tubulin (1:500, Sigma-Aldrich clone 6-11B-1), mouse monoclonal MUC5AC (1:250, Abnova clone 45M1).

Secondary antibodies: Alexa Fluor 488 goat anti-rabbit (1:500; Thermo Fisher Scientific), Alexa Fluor 647 goat anti-mouse (1:500; Thermo Fisher Scientific), Alexa Fluor 488 Goat anti-Mouse IgG1 (1:500, Fisher Scientific), Alexa Fluor 594 Goat anti-Mouse IgG2a (1:500, Fisher Scientific), Alexa Fluor 647 Goat anti-Mouse IgG2b (1:500, Fisher Scientific). Incubation with secondary antibodies was carried out during 1h at room temperature. Nuclei were stained with 4,6-diamidino-2-phenylindole (DAPI).

When necessary, Acetylated Tubulin, Muc5AC and KRT5 antibodies were directly coupled to CF 594/488/488 respectively, with the Mix-n-Stain kit (Sigma-Aldrich) according to the manufacturer’s instruction. Coupled primary antibodies were applied for 2 hours at room temperature after secondary antibodies had been extensively washed and after a 30 min blocking stage in 3% normal rabbit or mouse serum in PBS. MTECs immunostainings were directly performed on Transwell^®^ membranes using a similar protocol. For mounting on slides, Transwell^®^ membranes were cut with a razor blade and mounted with ProLong^TM^ Gold medium (Thermo Fisher Scientific). Images were acquired using the Olympus Fv10i or Leica sp5 confocal imaging systems.

### Time course sample analysis

#### Preprocessing

For each sample, cells with levels in the top 5% or bottom 5% of distribution for the following quality metrics: number of expressed features, dropout percentage and library size (total UMI count) were filtered out. Additionally, cells with a percentage of mitochondrial genes > top 5% were also removed. Quality metrics were computed using the scater package (2.3.0) [65]. Genes expressed at less than 1 UMI in at least 5 cells were removed from further analysis.

#### Normalization

The scran package [66] was used to calculate cell-based scale factors and normalize cells for differences in count distribution. Each sample was normalized separately twice, first in an unsupervised manner, then after grouping cells of similar gene expression based on our robust clustering results.

#### Clustering Robustness

In order to best determine the key steps in the studied differentiation process, a customized method was implemented to analyze clustering robustness to dataset perturbation. For all possible number of clusters (from 2 to 9), multiple subsets of the studied dataset were created (10 subsets, with 10% of the cells randomly removed each time) and clustering was performed multiple times on each subset with changing settings of the seed parameter. The result of those clusterings were stored in a n cells x n cells stability matrix, containing for each pair of cells 1 or 0 if the cells are clustered together or not. This stability matrix was then transformed in a Euclidean distance matrix between cells and then divided into the used k number of clusters k using hierarchical clustering (hclust with ‘average’ method). To identify the optimal number of clusters, a visual inspection of the elbow plot of the average intra-stability (mean stability within each cluster) and the average inter-stability (mean stability between each cluster) was done. Cells with a stability metric less than 70% were labelled as Unassigned, due to their high clustering variability between each round of clustering, then removed from further analysis of the time course data. Cell clustering was performed using SIMLR (package version 1.4.1) [67].

#### Differential analysis

To further analyze the robustness of each step of the differentiation process, we tested the robustness of the cell type marker gene identification through differential gene expression analysis. Differential expression analysis was performed using edgeR (package version 3.22) [68]. In a one vs. all differential analysis, a pool of 100 cells from one cluster were analyzed against an equal mixture of cells from all other clusters. In a one vs one differential analysis pools of cells of the same size were compared. Those differential analysis were performed multiple times (10 times) on different pool of cells and the DEG identified were compared between each pool of cells using the rank-rank hypergeometric overlap algorithm [69]. Unfortunately, this approach was too stringent, and only identified highly expressed marker gene as they are less likely to submit to dropout events. Thus, the Seurat FindAllMarkers function based on a non-parametric Wilcoxon rank sum test was used to identify cell type marker genes.

#### Time points aggregation

10X datasets generated during the time course were aggregated using MNN correction [70] from the scran package.

#### Trajectory Inference

Trajectory inference was performed using monocle 2 (package version 2.8) [71]. Cell ordering was based on highly variable genes (~ 200 −500 genes) selected by their expression dispersion. Monocle analysis on the aggregated time points was done on raw count after library size correction (downsampling). Branch building was performed using BEAM analysis within the monocle package, and corresponding differential analysis was done by cross comparison of group of cells along the pseudotime (before branching, after branching and at the branch end) using Seurat 1 vs 1 differential analysis.

#### Cell type projection

To compare cell types identified in distinct samples, cells were projected from one dataset onto the other using scmap R package version 1.1, scmapCluster function [72].

#### Data Visualization

All graphs were generated using R (ggplot2). Heatmaps were obtained using pheatmap (no clustering used, genes ordered by their expression in pseudotime or in cluster, cells ordered by pseudotime or cluster). Heatmaps show smoothed gene expression values: for each gene, normalized gene expression values were first transformed into z-scores, then averaged across 10 neighboring cells in the chosen ordering (pseudotime only or pseudotime in clusters). For single gene representation: only cells with expression levels above the top 50 percentiles for that gene are represented for clarity.

### Individual sample analysis

Each sample of our study was reanalyzed with less stringent parameters, to identify rare or transitory cell types or gene expression events.

#### Preprocessing, normalization and clustering

Individual dataset analysis was performed using Seurat standard analysis pipeline. Briefly, cells were first filtered based on number of expressed features, dropout percentage, library size and mitochondrial gene percentage. Thresholds were selected by visually inspecting violin plots in order to remove the most extreme outliers. Genes expressing less than 5 UMI across all cells were removed from further analysis. Cell-level normalization was performed using the median UMI counts as scaling factor. Highly variable genes were selected for following analysis based on their expression level and variance. PCA analysis was performed on those hvgs, the number of PCs to use was chosen upon visual inspection of the PC variance elbowplot (~10 to 20 PCs depending on the dataset). Clustering was first performed with default parameter and then increasing the resolution parameter above 0.5 to identify small clusters (but with the knowledgeable risk of splitting big cluster due to high gene expression variability). Differential analysis was again performed using Seurat FindAllMarkers and FindMarkers functions based on non-parametric Wilcoxon rank sum test. Gene Set Enrichment analysis was performed using fgsea R package with the following gene sets reactome.db (R package) and GO cellular component (Broad Institute GSEA MSigDB) genesets. Molecular function enrichment analysis was performed using Ingenuity Pathway Analysis (Qiagen).

#### Cell Type Annotation

Based on the time course experiment analysis and associated top ~15 marker genes identified, a score was computed to associate cell types to each cluster. Scoring method is based on Macosko*. et al.* cell cycle phase assignment [73]. For each cell it measures the mean expression of the top marker genes for each possible cell type, which results in a matrix c cell types per n cells. Then it performs a z-score of those mean expression for each cell, the top resulting score gives the matching cell type.

#### Velocity

RNA velocity was calculate using latest release of velocyto pipeline (http://velocyto.org/) using standard parameters: GTF file used for Cell Ranger analysis and the possorted_genome_bam.bam, Cell Ranger output alignment file. From the loom file which contains count table of spliced and unspliced transcript the gene.relative.velocity.estimates function was used on cell type marker genes. The resulting expression pattern of unspliced-spliced phase portraits shows the induction or repression of those marker genes from one cell type to the next. We used velocyto package version 0.5 [33].

### Plasscheart *et al.* dataset

Plasscheart *et al.* data [29] were downloaded as processed data along with visualization coordinates and were used without further manipulation. (https://kleintools.hms.harvard.edu/tools/springViewer_1_6_dev.html?datasets/reference_HBECs/reference_HBECs)

## Declarations

### Ethics approval and consent to participate

Human samples were collected according to the guidelines of the Declaration of Helsinki, after approval by the institutional review board “Comité de Protection des Personnes Sud Est IV” (12/12/2017). All patients gave their written informed consent to participate. Mouse experiments were performed following the Directive 2010/63/EU of the European parliament and of the council of 22 September 2010 on the protection of animals used for scientific purposes. The IPMC is authorized to use animals by the French agricultural ministry (Ministere de l’Agriculture et de l’Alimentation) and persons performing experiments were authorized by the French research ministry (Ministère de l’Education Supérieure, de la Recherche et de l’Innovation).

### Consent for publication

All patients gave their written informed consent for anonymous publication of data arising from their samples.

### Availability of data and materials

All datasets generated during this study were deposited to the Gene Expression Omnibus repository under the series number GSE121600.

### Authors’ contributions

LEZ and PB designed and supervised the study and obtained fundings. SRG, MJA, AC, VM and LEZ performed the experiments. SL et CHM provided human fresh tissues. IC provided pig trachea. MD, KLB and AP performed the bioinformatics analysis. All authors were involved in data interpretation. MD and LEZ designed the figures. LEZ drafted the original manuscript. MD, PB, SRG, AP, KLB, BM and LEZ edited the manuscript.

## Acknowledgements

We are grateful to the UCAGenomiX platform and to Marin Truchi for fruitful discussions and technical help on single cell RNA sequencing.

## Supplementary figures

**Supplementary figure 1:**
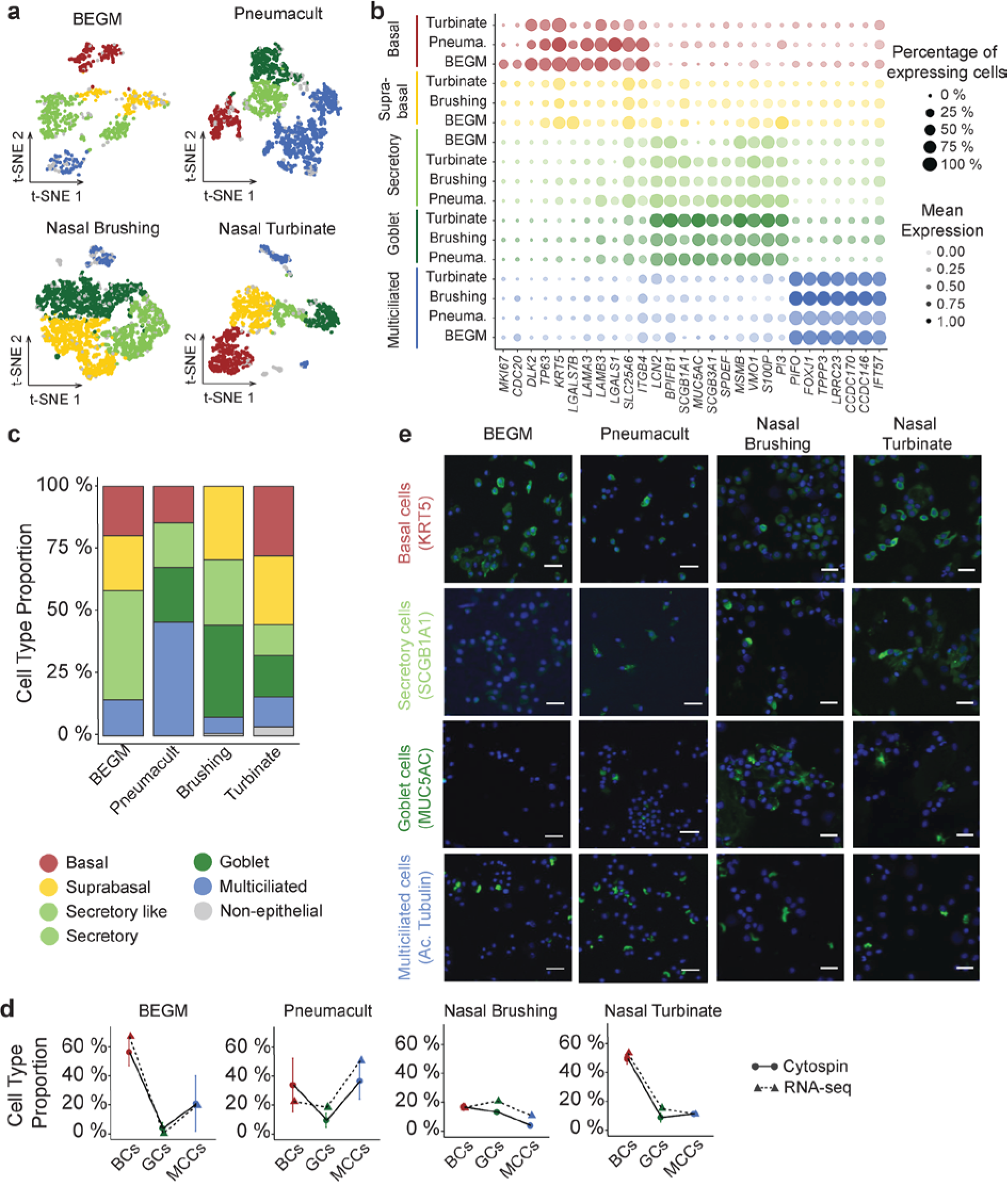
Cell type composition comparison between homeostatic *in vitro* samples and fresh human tissues. **a** t-SNE representing the distinct cell populations identified in each sample. **b** Dot plot of the main cell population marker genes. Dot size describe the percentage of cells expressing the respective marker genes and the average expression level of that gene based on UMI counts are shown by color intensity. **c** Relative abundance of cell types in each sample. **d** Scatter plot of the percentage of cells expressing the selected marker genes for basal, goblet and multiciliated cells at the transcript (dotted line) and protein level (full line). **e** Immunostaining of the selected marker genes for basal, secretory, goblet and multiciliated in cytospin for each sample.

**Supplementary figure 2:**
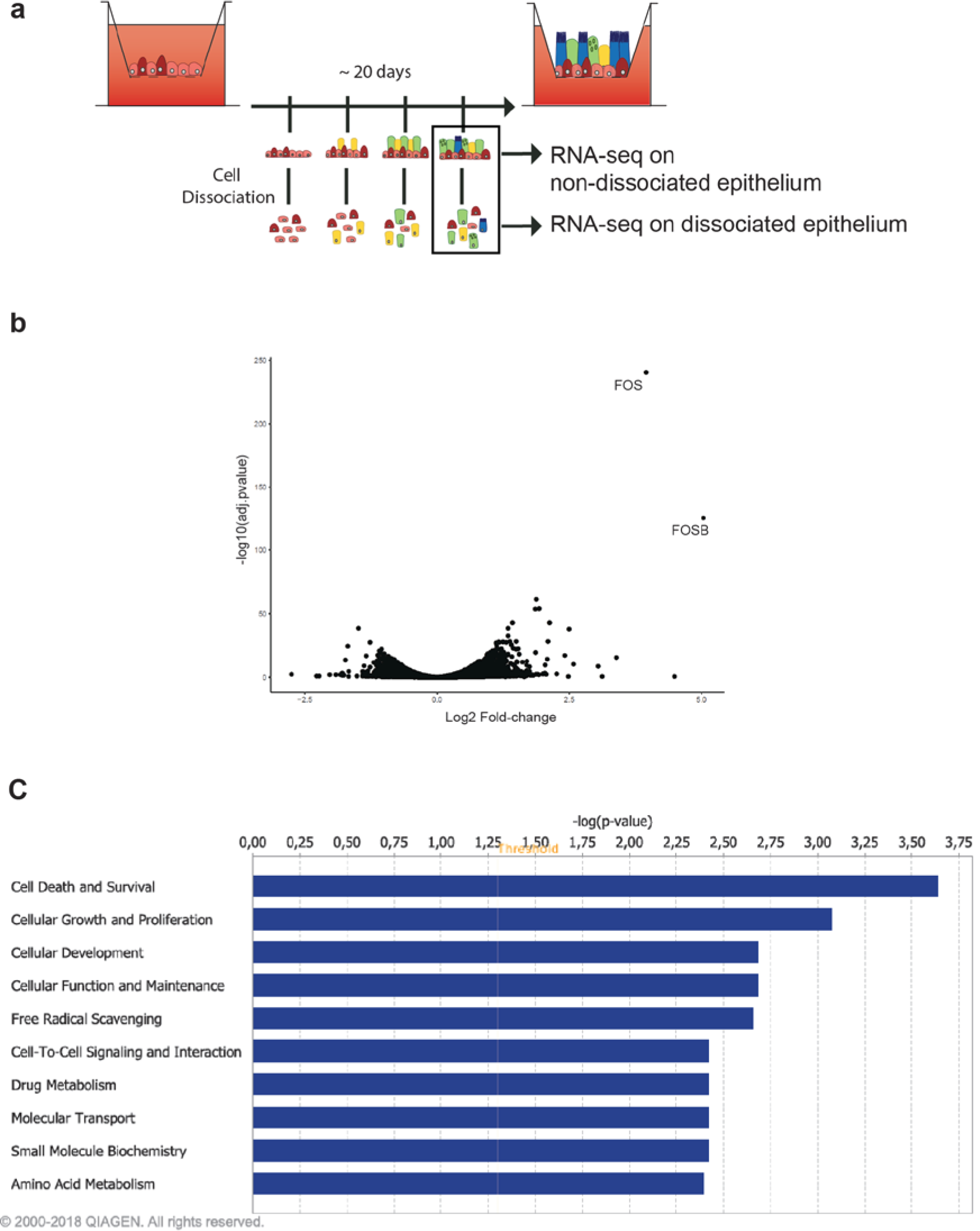
Effect of cell dissociation on HAEC gene expression. **a** RNA-seq experimental design. Regenerating airway epithelia was dissociated at 28 days after a transition to an air-liquid interface. Dissociated and non-dissociated cultures from 2 donors were subjected to RNA-seq. **b** Volcano plot showing differential gene expression of dissociated vs. non-dissociated cell cultures. **c** Identification, with Ingenuity Pathway Analysis (Qiagen) of the molecular functions most significantly affected by dissociation, based on an analysis of 300 differential expressed genes (FDR<0.01 and abs(log2FC)>1).

**Supplementary figure 3:**
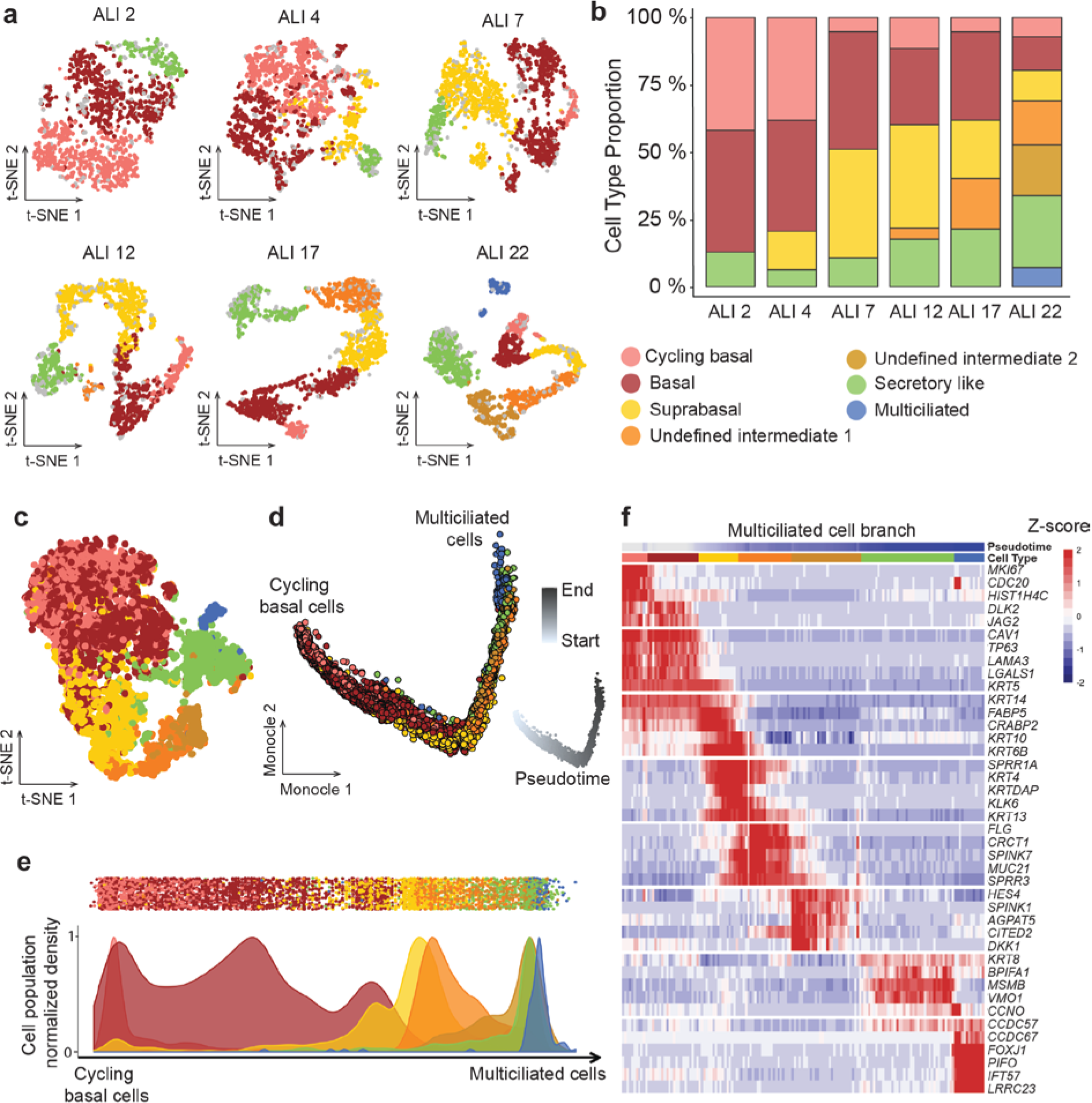
Characterization of MCC cell lineages during airway epithelium regeneration using single cell RNA-Seq in BEGM medium. **a** t-SNE representing the distinct cell populations identified at each time points. **b** Relative abundance of cell types at each time point. **c** t-SNE plot of the aggregate of all cells from each time point. **d** Representation of the cell lineages inferred by Monocle 2 occurring during the upper airway epithelium regeneration (aggregate of all time points). Pseudotime evolution along the differentiation trajectory shown by white to grey gradient. **e** Distribution of the defined cell types in pseudotime. **f** Heatmap representing the smoothed temporal expression pattern of indicated cell type specific marker genes on MCC trajectory. Cells were ordered by cluster appearance in pseudotime.

**Supplementary figure 4:**
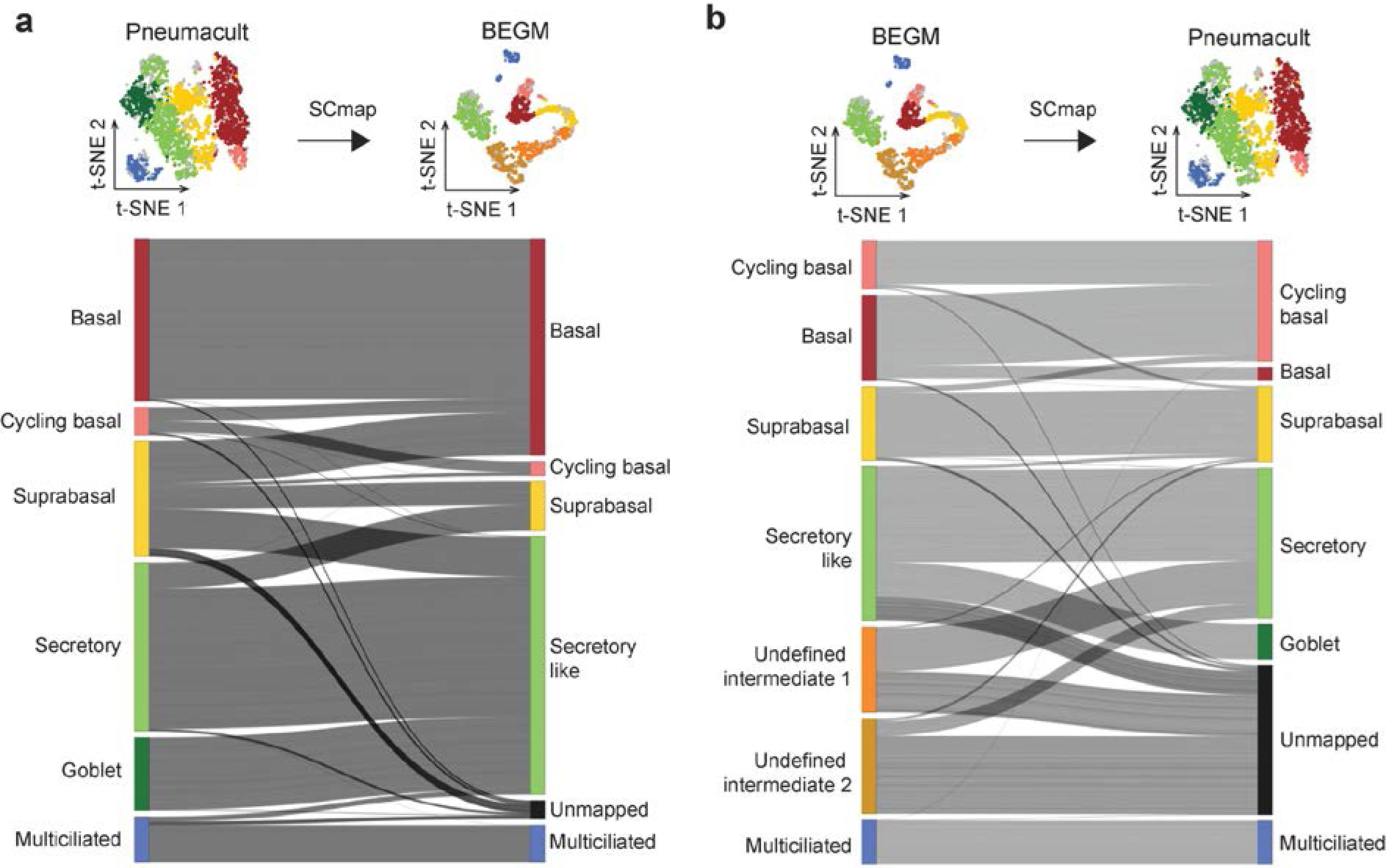
Comparison of fully differentiated epithelia cell population between Pneumacult and BEGM media. **a** Sankey Network of the mapping of Pneumacult cells onto BEGM cells. **b** Sankey Network of the mapping of BEGM cells onto Pneumacult cells.

**Supplementary figure 5:**
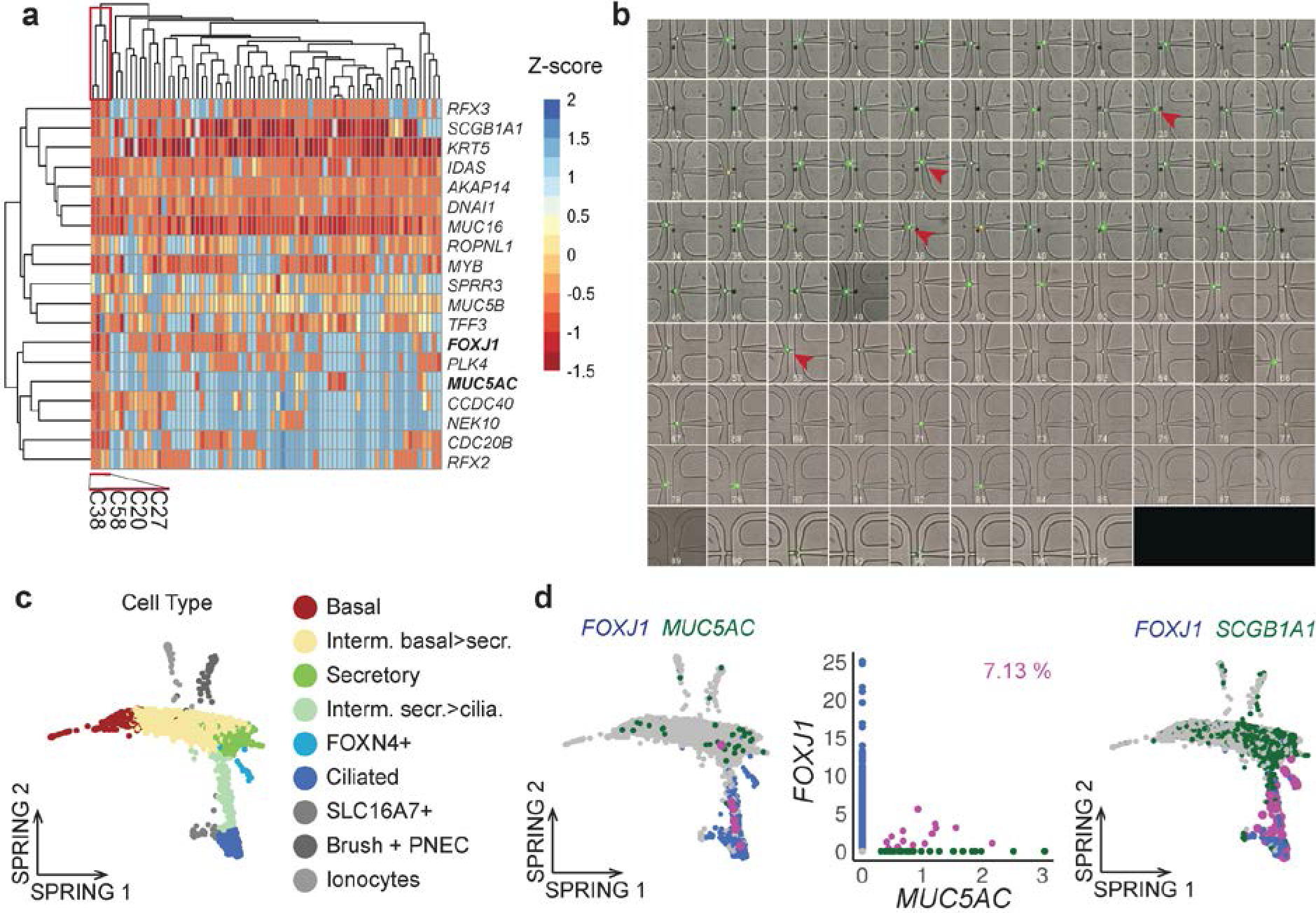
Validation of Goblet cells markers in multiciliated cells in doublet free single cell transcriptomic datasets. **a** Heatmap colored by Z-score from C1 Biomark experiment in PneumaCult media, co-expression of Goblet and Multiciliated cell markers in single cells (red cluster). **b** C1™ Single-Cell Preamp IFC (10–17 µm) imaging, Red arrowheads show cells expressing both cell type markers (*FOXJ1* and *MUC5AC*), one cell per chamber, green cells=living cells, red cells=dead cells. **c** SPRING representations of Plasschaert *et al.* dataset. Left panel colored by cell type. Center left panel colored by *FOXJ1* + cells (blue), *MUC5AC* + cells (green), co-expressing *FOXJ1* and *MUC5AC* cells (pink). Center right panel displays a scatter plot of normalized expression of *MUC5AC* and *FOXJ1* in cells (dot) colored by *FOXJ1* + cells (blue), *MUC5AC* + cells (green), co-expressing FOXJ1 and MUC5AC cells (pink). Right panel colored by *FOXJ1* + cells (blue), *SCGB1A1* + cells (green), co-expressing *FOXJ1* and *SCGB1A1* cells (pink).

**Supplementary figure 6:**
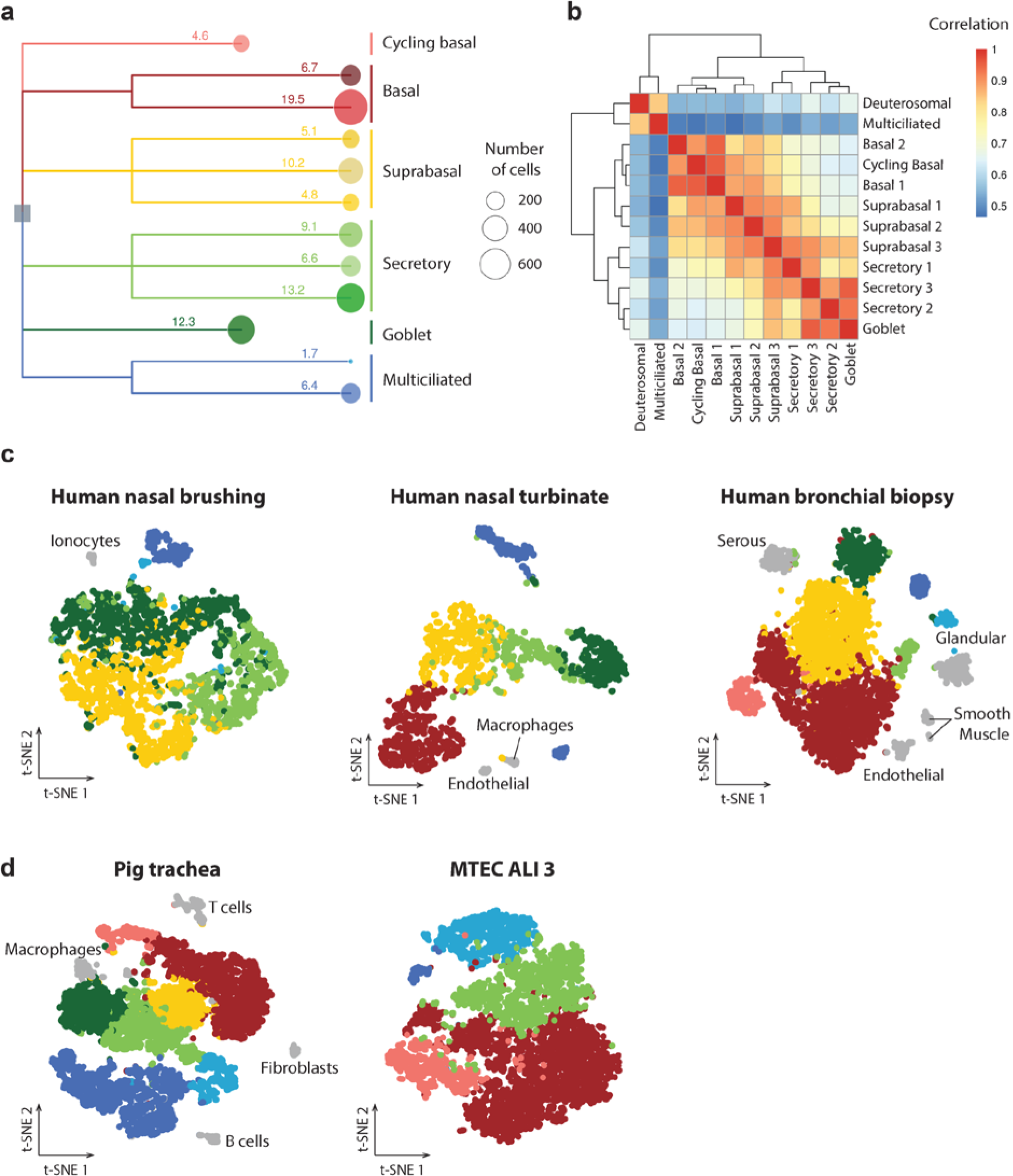
Identification of subpopulations by high resolution cell clustering. **a** Dendrogram of cell types identified through robust and high-resolution clustering of scRNA-seq data from Pneumacult ALI28 sample. **b** Correlation matrix of cell type identified in Pneumacult sample. **c** tSNE representation of scRNA-seq data from human nasal brushing (left), nasal turbinate (center), bronchial biopsy (right) colored by cell type described previously (Fig. 1), in grey are rare or nonepithelial cell types. **d** t-SNE representation of scRNA-seq data from pig trachea (left) and mouse culture (right) colored by cell type previously described (Fig. 1).

**Supplementary figure 7:**
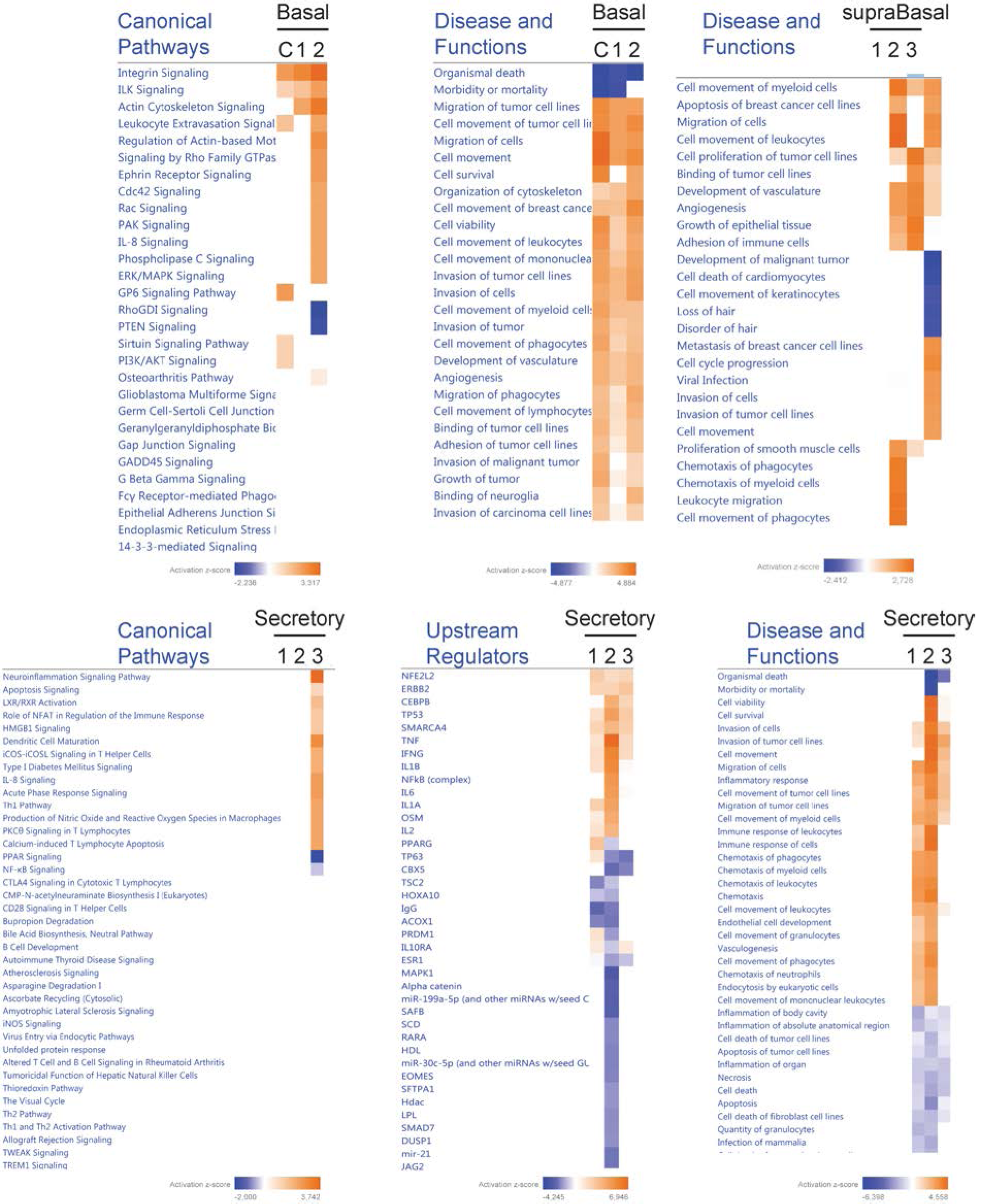
Pathway enrichment comparison between basal, suprabasal and secretory cell subtypes. Identification, with Ingenuity Pathway Analysis (Qiagen) of the enriched canonical pathways, upstream regulators and disease and functions in each of the basal and secretory subpopulations, based on an analysis of 300 differential expressed genes (FDR<0.01 and log2FC>0.6 for BCs and SCs, and log2FC>0.4 for supraBCs). C: cycling basal cells.

**Supplementary figure 8:**
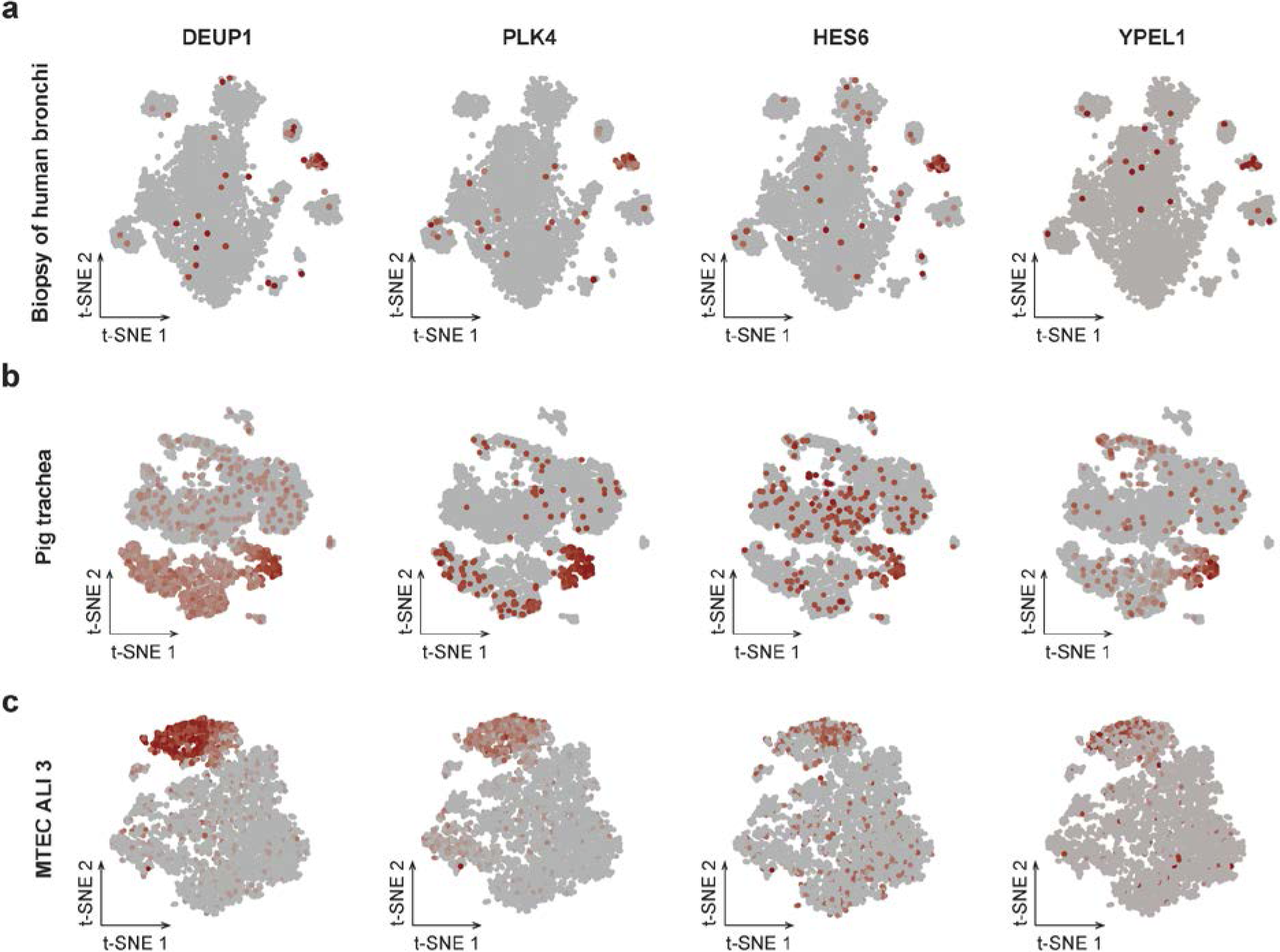
Robustness of Deuterosomal cells marker genes. **a, b, c** t-SNE representation of Deuterosomal cell population marker genes expression (lowly to highly expressed, grey to red) in (a) Biopsy of human trachea, (b) Pig trachea, (c) MTECs ALI 3.

**Supplementary figure 9:**
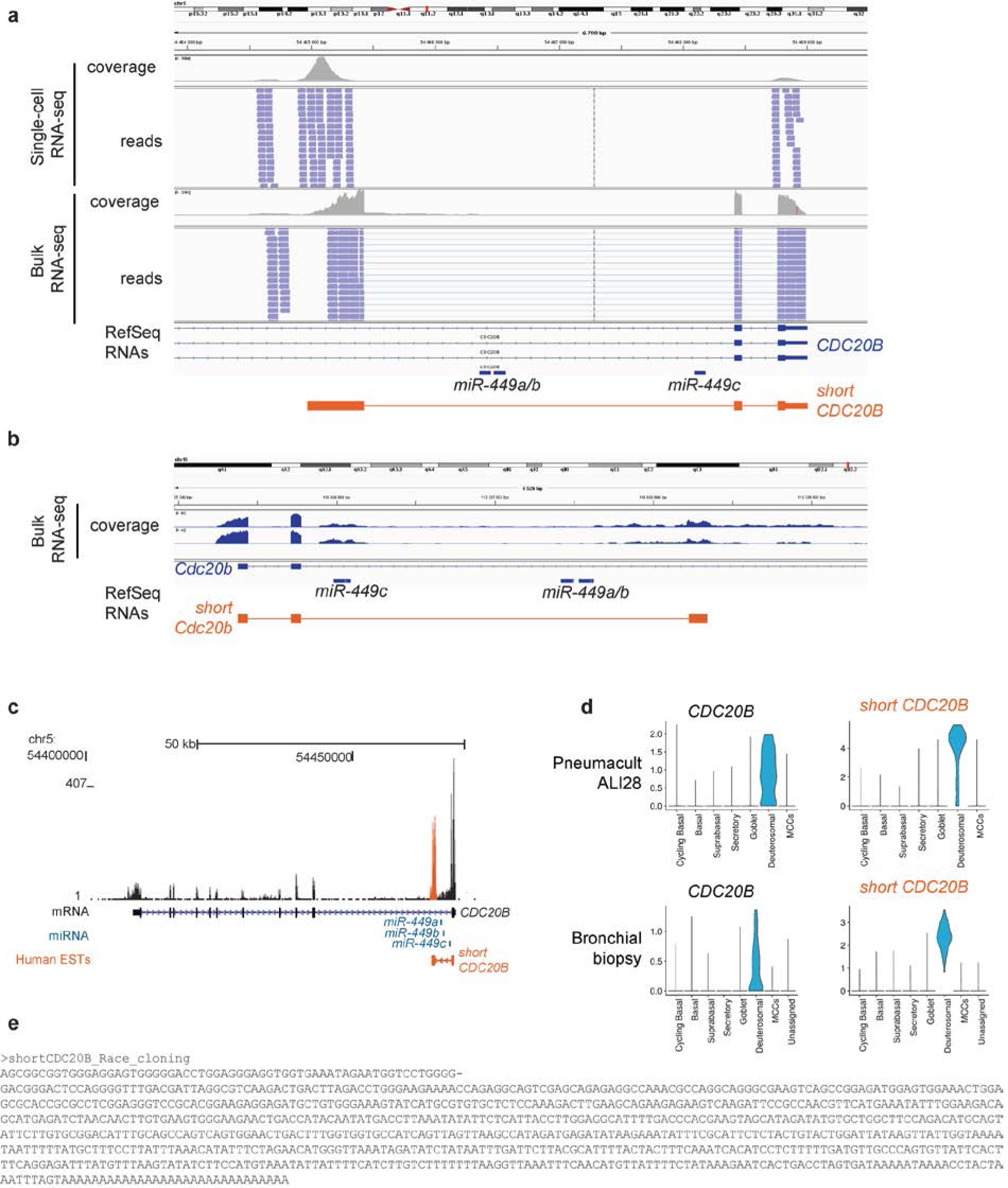
Identification of short CDC20B, a novel isoform of a deuterosomal cell population marker gene. **a** Integrated Genome Viewer (hg19) view of the CDC20B gene with coverage and read alignment from bulk RNA-seq of well-differentiated HAECs and single-cell RNAseq of ALI14 differentiated HAECs. **b** Genome Viewer (mm10) view of the CDC20B gene with coverage from bulk RNA-seq of ALI 7 differentiated MTECs from the public dataset GSE75715. **c** UCSC Genome browser (hg19) view of the CDC20B gene with coverage from read alignment from bulk RNA-seq of ALI28 differentiated HAECs. The alternative 3rd exon is shown in orange. **d** Violin plots for CDC20B and short CDC20B abundance in the Pneumacult ALI28 and the bronchial biopsies samples. **e** Alternative human short CDC20B sequence identified with 5’ race cloning.

**Supplementary figure 10:**
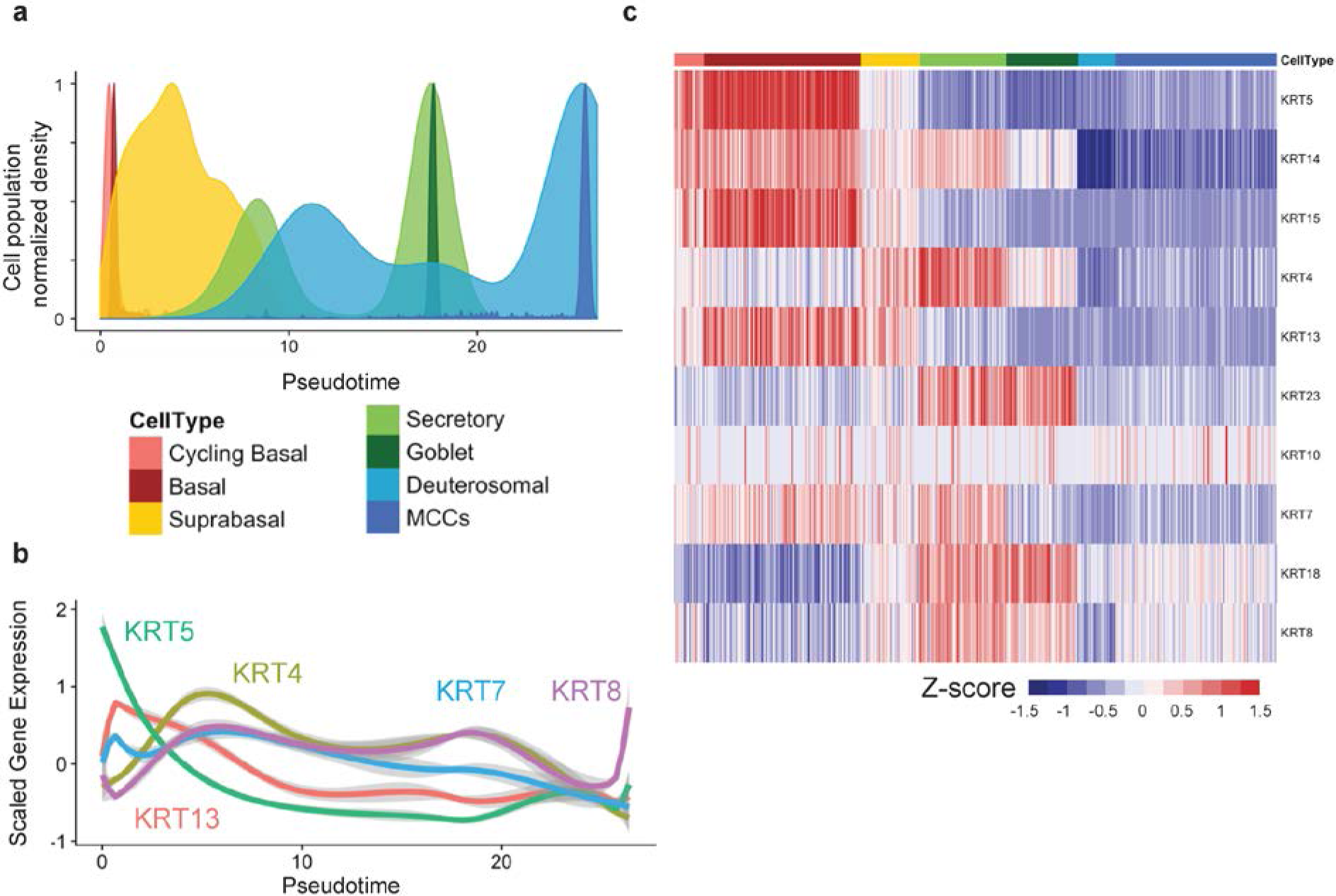
Keratin expression in scRNA-seq from pig tracheal cells. **a** Distribution of the 7 main cell types in the pseudotime from scRNA-seq of pig tracheal cells. **b** Plot of normalized gene expression of selected keratins according to pseudotime. **c** Heatmap representing the smoothened temporal expression pattern of regulated keratins. Cells were ordered by cluster appearance.

**Supplementary figure 11:**
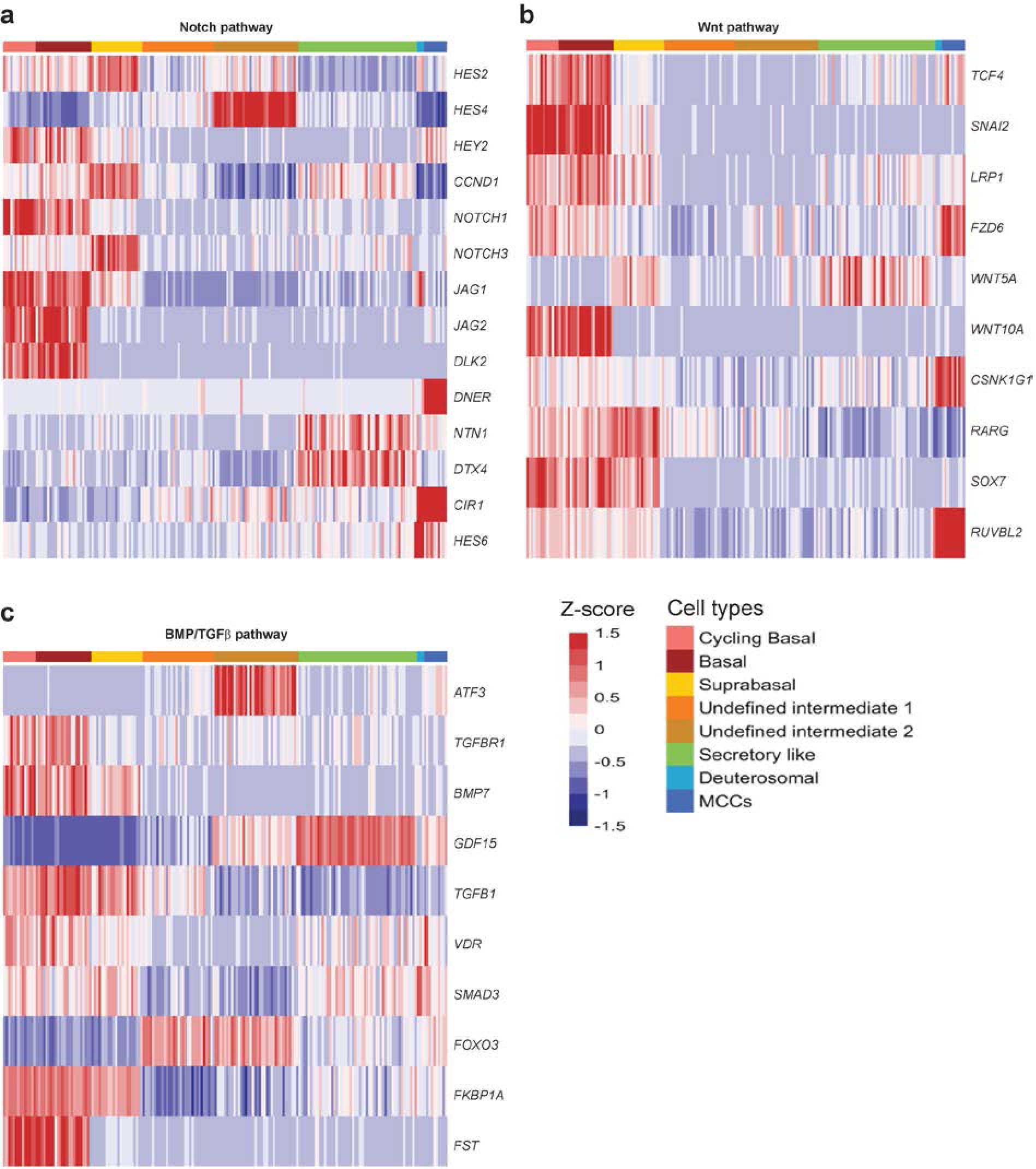
Single-cell expression of signaling pathways components during airway regeneration in BEGM medium. **a** Heatmap of the genes related to the NOTCH pathway with cells ordered by cluster. **b** Heatmap of the genes related to the WNT pathway with cells ordered by cluster. **c** Heatmap of the genes related to the BMP/TGFβ pathway with cells ordered by cluster.

## Supplementary tables

**Supplementary table 1:**
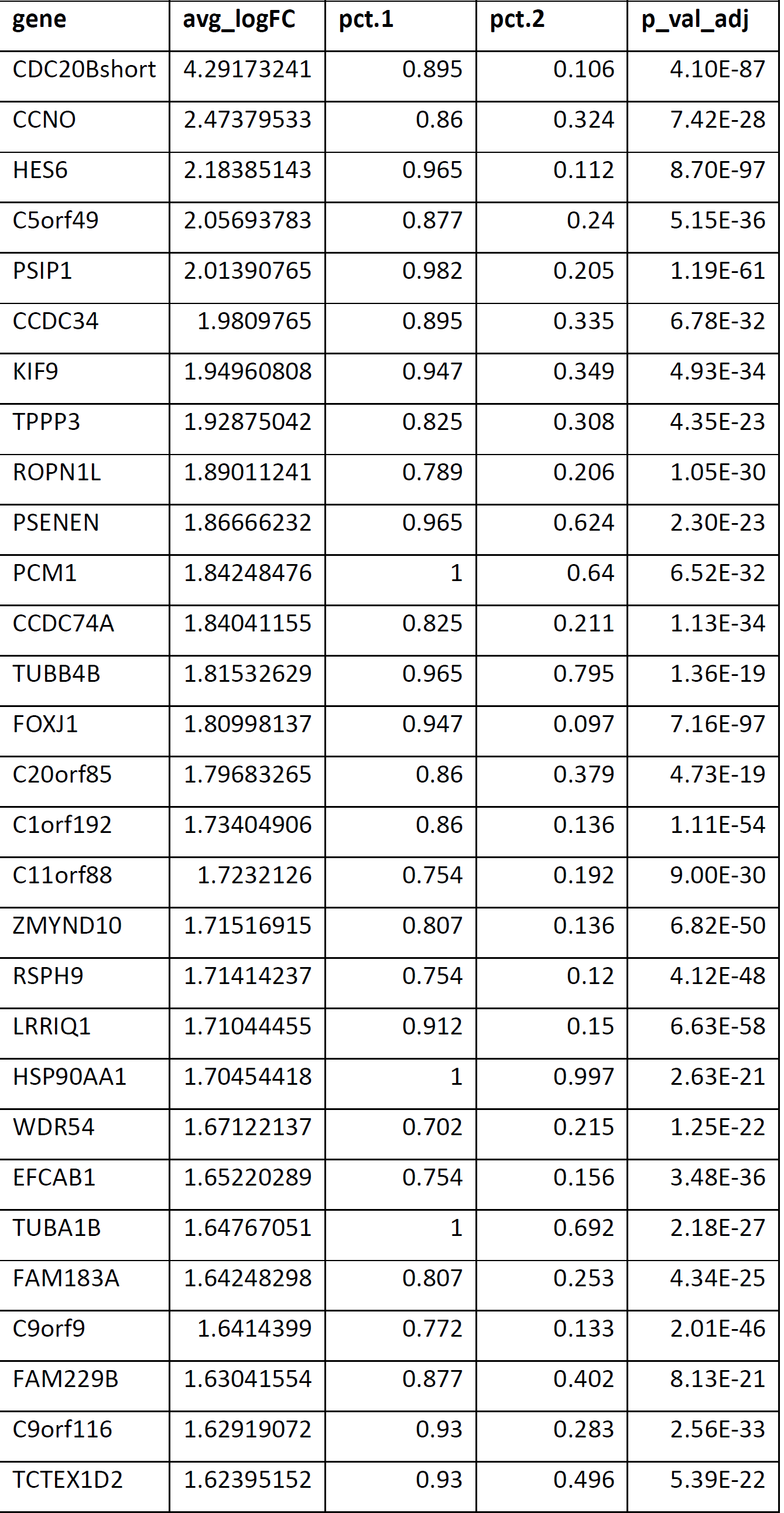
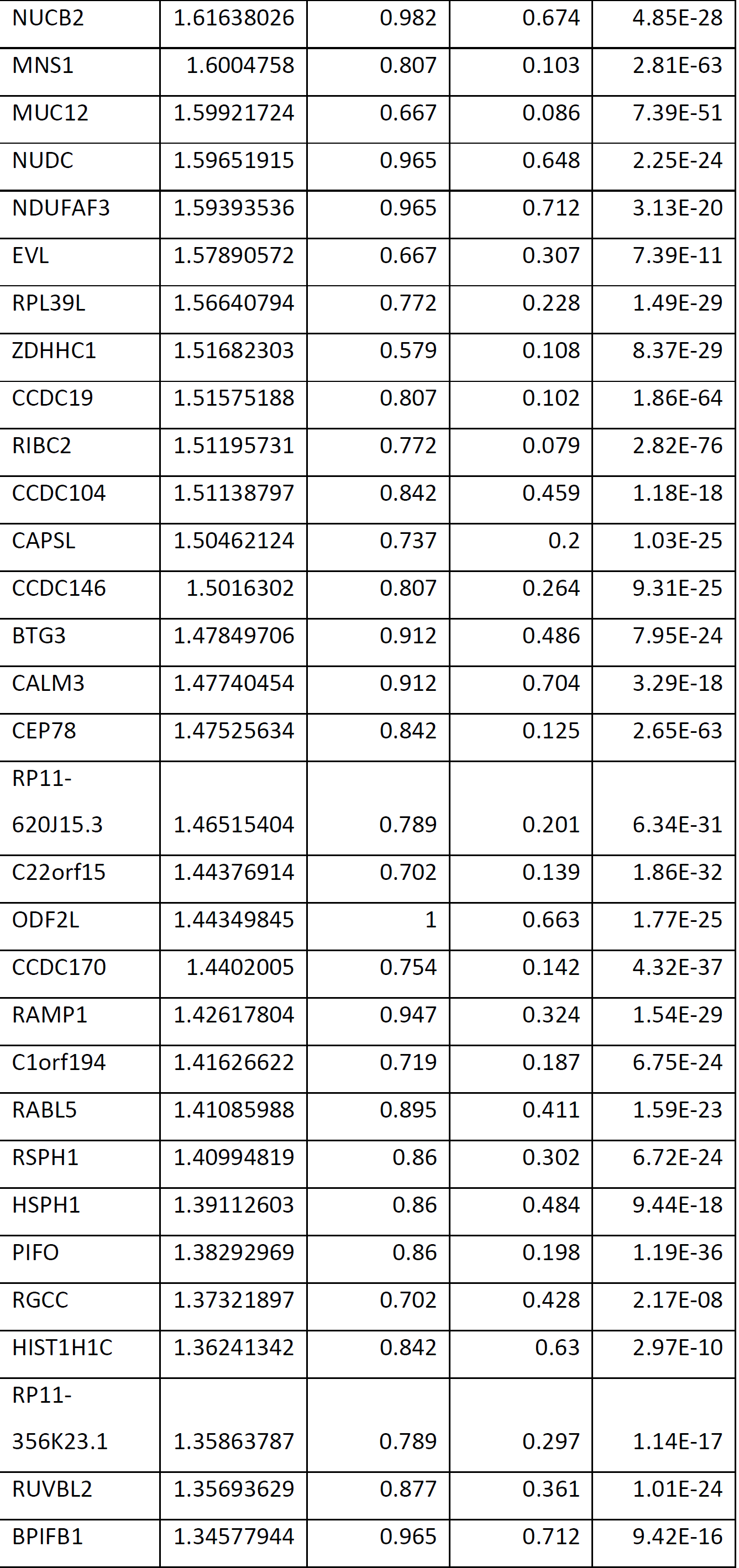
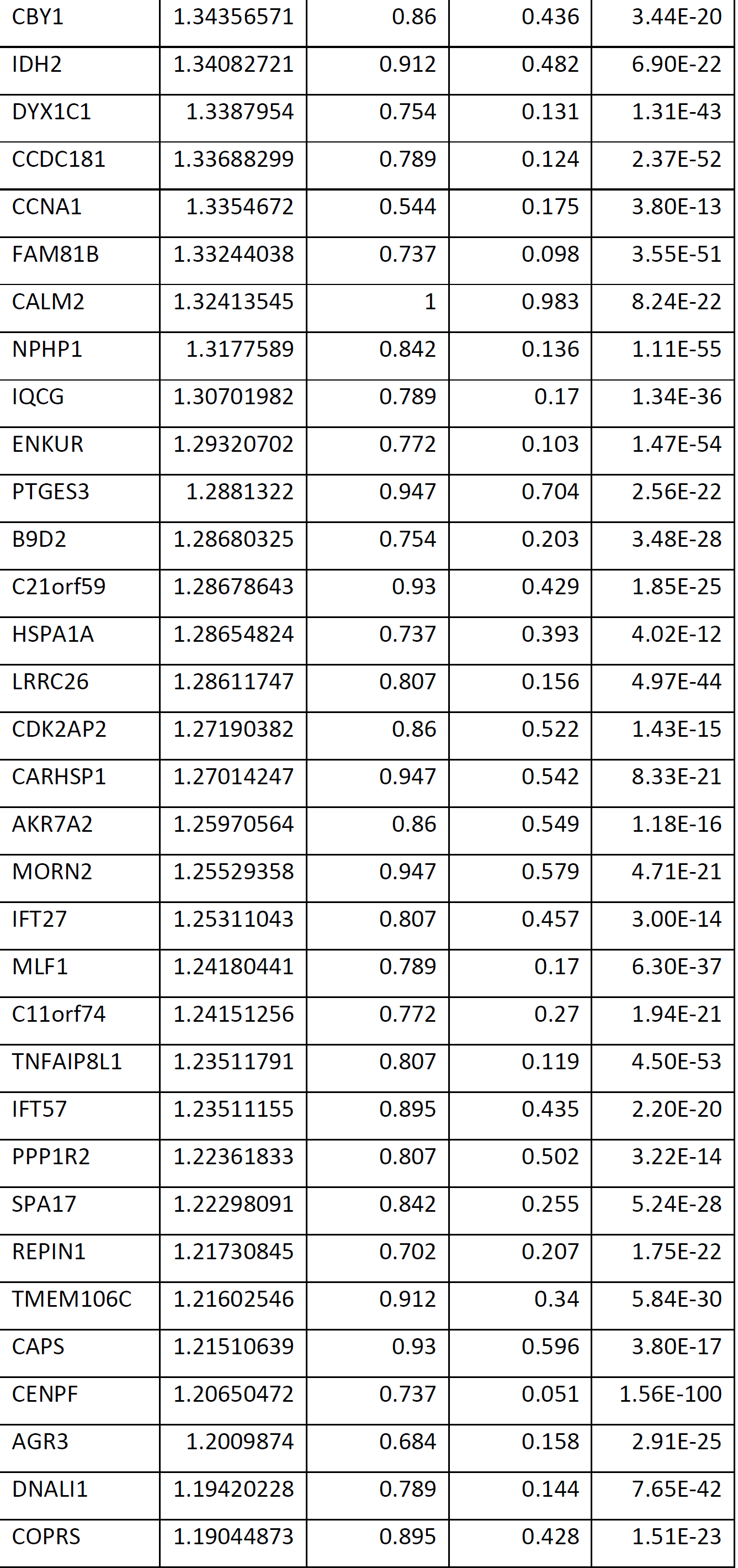
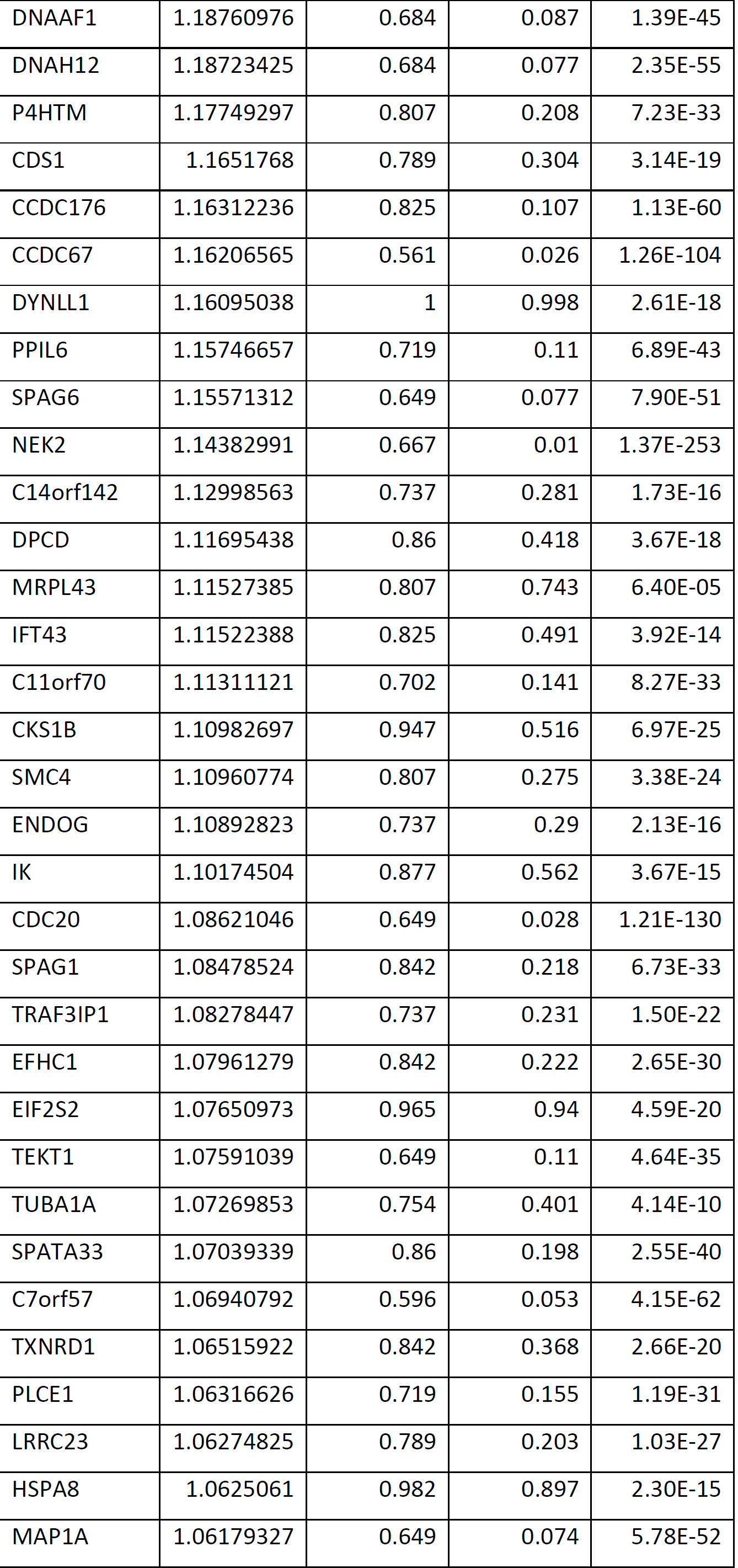
Differential Gene expression of Deuterosomal cluster vs. All clusters.

**Supplementary table 2:**
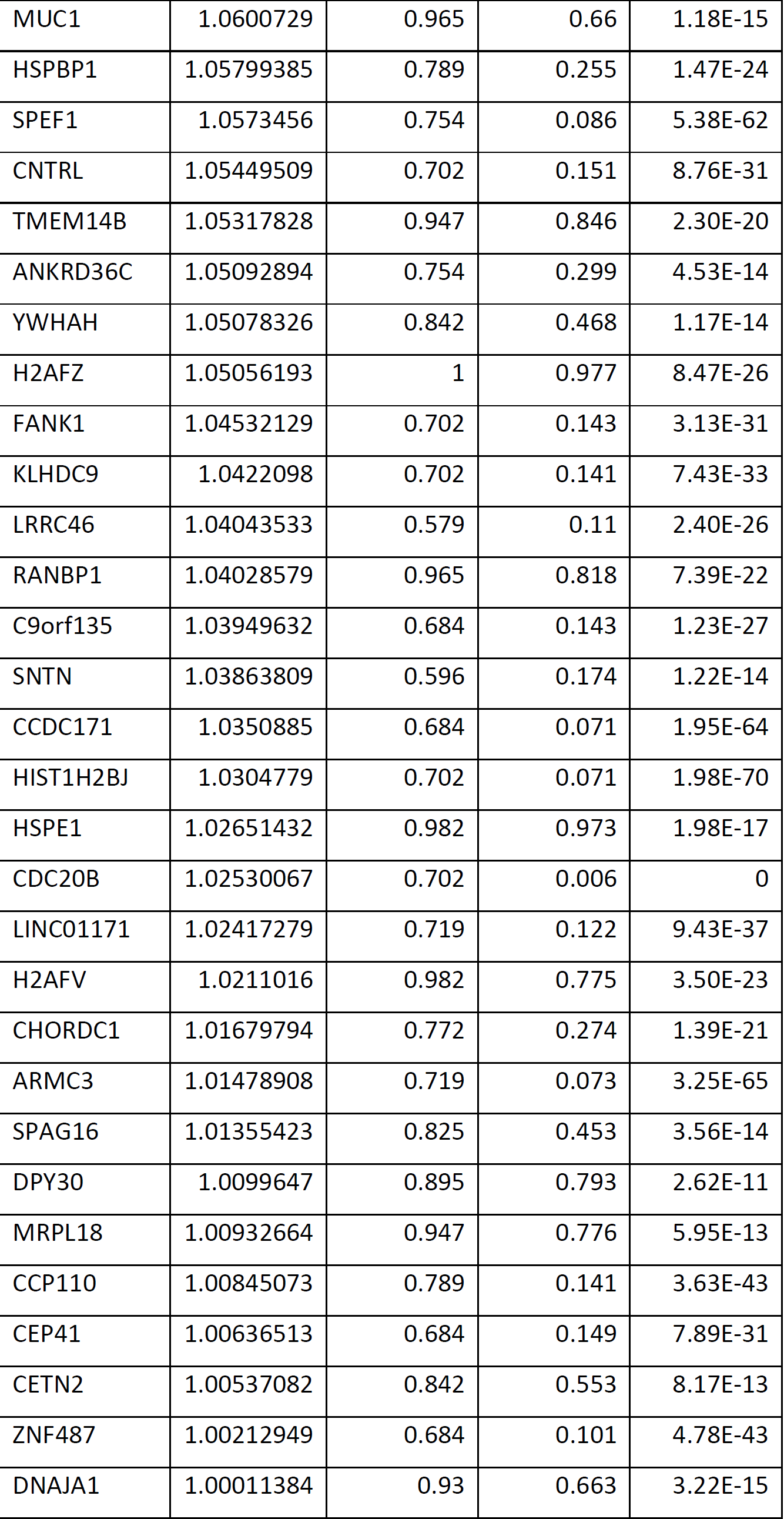
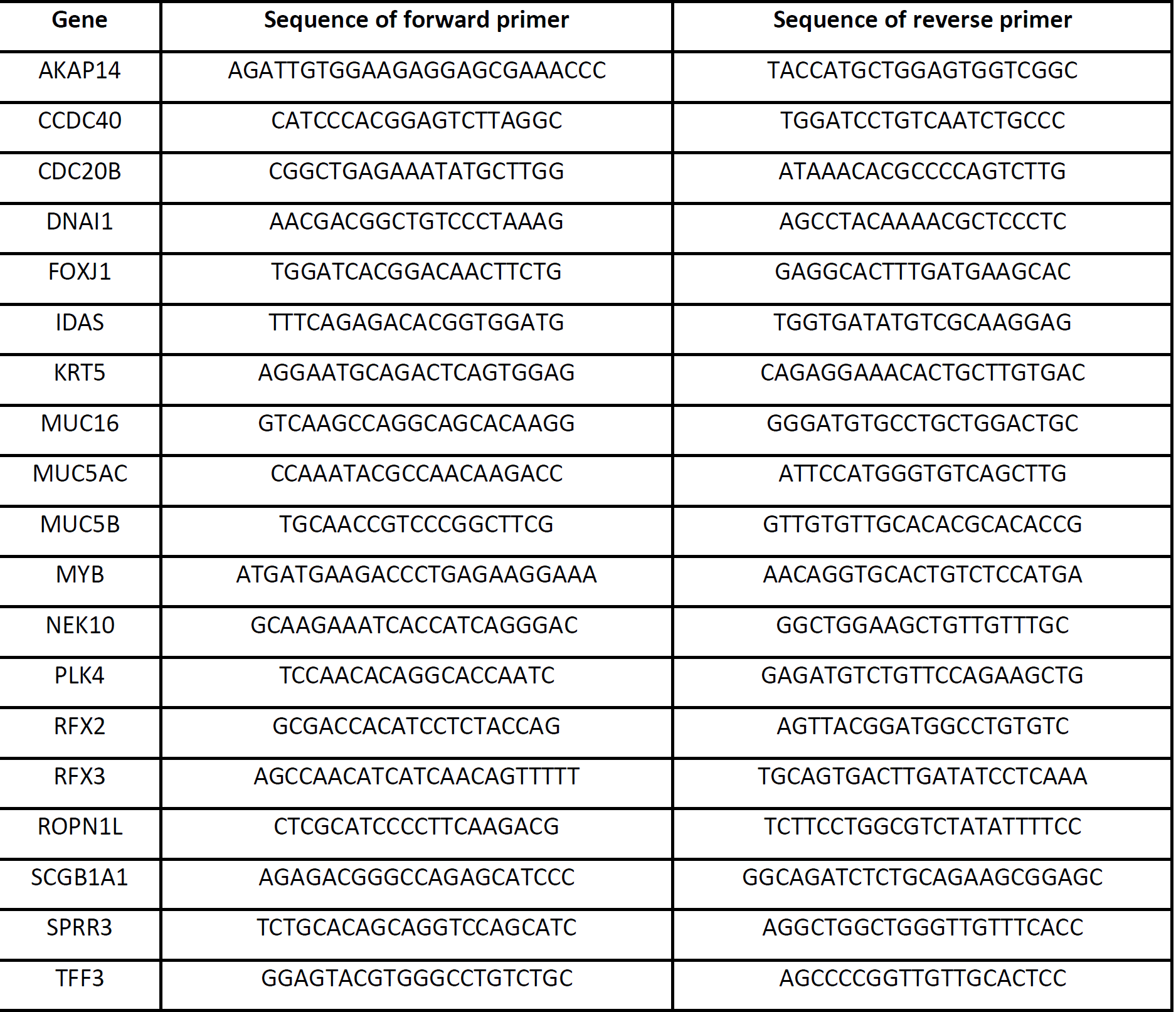
Primers used in the Biomark qRT-PCR.

